# A supervised Bayesian factor model for the identification of multi-omics signatures

**DOI:** 10.1101/2023.01.25.525545

**Authors:** Jeremy P. Gygi, Anna Konstorum, Shrikant Pawar, Edel Aron, Steven H. Kleinstein, Leying Guan

## Abstract

**Motivation:** Predictive biological signatures provide utility as biomarkers for disease diagnosis and prognosis, as well as prediction of responses to vaccination or therapy. These signatures are identified from high-throughput profiling assays through a combination of dimensionality reduction and machine learning techniques. The genes, proteins, metabolites, and other biological analytes that compose signatures also generate hypotheses on the underlying mechanisms driving biological responses, thus improving biological understanding. Dimensionality reduction is a critical step in signature discovery to address the large number of analytes in omics datasets, especially for multi-omics profiling studies with tens of thousands of measurements. Latent factor models, which can account for the structural heterogeneity across diverse assays, effectively integrate multi-omics data and reduce dimensionality to a small number of factors that capture correlations and associations among measurements. These factors provide biologically interpretable features for predictive modeling. However, multi-omics integration and predictive modeling are generally performed independently in sequential steps, leading to suboptimal factor construction. Combining these steps can yield better multi-omics signatures that are more predictive while still being biologically meaningful.

**Results:** We developed a supervised variational Bayesian factor model that extracts multi-omics signatures from high-throughput profiling datasets that can span multiple data types. Signature-based multiPle-omics intEgration via lAtent factoRs (SPEAR) adaptively determines factor rank, emphasis on factor structure, data relevance and feature sparsity. The method improves the reconstruction of underlying factors in synthetic examples and prediction accuracy of COVID-19 severity and breast cancer tumor subtypes.

**Availability:** SPEAR is a publicly available R-package hosted at https://bitbucket.org/kleinstein/SPEAR.

## 1 Introduction

Biological signatures, composed of subsets of predictive analytes, serve as valuable biomarkers for disease diagnosis and prognosis (Ramilo *et al*., 2007; Yu *et al*., 2019; Bodkin *et al*., 2022; Chawla *et al*., 2022), as well as the prediction of responses to vaccinations and therapies (Fourati *et al*., 2022; Hagan *et al*., 2022). In addition to being predictive, signatures offer interpretable insight into the underlying mechanisms that drive disease, further adding to their value to the scientific community. By leveraging associations between different types of analytes, multi-omics signatures offer the possibility of improved performance and greater insights.

The identification of predictive biological signatures continues to be a prominent area of focus in biomedical research. For example, Fourati et al. (Fourati *et al*., 2022) and Hagan et al. (Hagan *et al*., 2022) identified pre- and post-vaccination gene signatures predictive of antibody responses across 13 different vaccines by random forest classification and logistic regression, respectively. Walker et al. (Walker *et al*., 2023) identified potential protein biomarkers of dementia via Cox proportional hazards regression modeling. Signatures can also provide mechanistic insight into underlying biological processes that affect the response of interest. Nakaya et al. (Nakaya *et al*., 2011) found a gene signature predictive of seasonal TIV vaccination response that included TLR5, known to sense bacterial flagellin. High association between TLR5 and TIV vaccination prompted the study and hypothesis that other ligands for TLR5, such as microbiota, are involved in influencing adaptive immunity to vaccination. These studies, like most studies, identify signatures that are derived from a single biological assay.

Recent advancements in high-throughput technologies have enabled the simultaneous collection of data from multiple assays (or ‘omics) from the same sample (Bhattacharya *et al*., 2014). These multi-omics datasets have the potential to yield multi-omics signatures that integrate biological signals across different modalities to predict a response of interest. While the dimensionality of multi-omics datasets is high, often only a small number of key underlying factors are used to capture the major data variation from the different assays. For example, activation of the interferon signaling pathway can drive hundreds of genes (Bolen *et al*., 2014), proteins (Lazear *et al*., 2019), and metabolites (Banoth and Cassel, 2018) to vary together across samples. One option to identify these major sources of variation is through the employment of latent factor models (Cantini *et al*., 2021), which construct low-dimensional factors from groups of biological analytes via dimensionality reduction. Unsupervised dimensionality reduction techniques for discovering factors are widely used in practice, including both non-probabilistic approaches such as principal component analysis (PCA), single and multi-block canonical correlation analysis (CCA) (Tenenhaus *et al*., 2014; Tenenhaus and Tenenhaus, 2011) as well as probabilistic approaches based on Bayesian factor models including Multi-omic Factor Analysis (MOFA) (Argelaguet *et al*., 2020) and iClusterBayes (Mo *et al*., 2018).

Conventionally, when identifying predictive signatures, the process of data integration and predictive modeling are performed separately, with multiomics factors being constructed before subsequent association with a response of interest. Combining these steps can lead to better multi-omics signatures that are more predictive while still being biologically interpretable (Singh *et al*., 2019; Li *et al*., 2012). To accomplish this, we present Signature-based multiPle-omics intEgration via lAtent factoRs (SPEAR). The SPEAR model employs a probabilistic Bayesian framework to jointly model multi-omics data with response(s) of interest, emphasizing the construction of predictive multi-omics factors. SPEAR estimates analyte significance per factor, extracting the top contributing analytes as a signature. In addition, the SPEAR model is amenable to various types of responses in both regression and classification tasks, permitting both continuous responses such as antibody titer and gene expression values, as well as categorical responses like disease subtypes. We demonstrate SPEAR’s advantages under simulated settings and validate SPEAR’s improved performance on real public multi-omics datasets for breast cancer and SARS-CoV-2 (COVID-19) through multi-class area under the receiver operating characteristic (AUROC) testing and balanced misclassification errors.

## 2 Methods

SPEAR identifies multi-omics signatures through the construction of predictive factors via dimensionality reduction. SPEAR jointly models both multiomics assays (*X*) and the response of interest (*Y*) and approximates posteriors of the factor loadings (*β*) and factor scores (*U*) using the variational Bayes inference (Fig. 1, Supplementary Methods B2).

**Figure 1.**
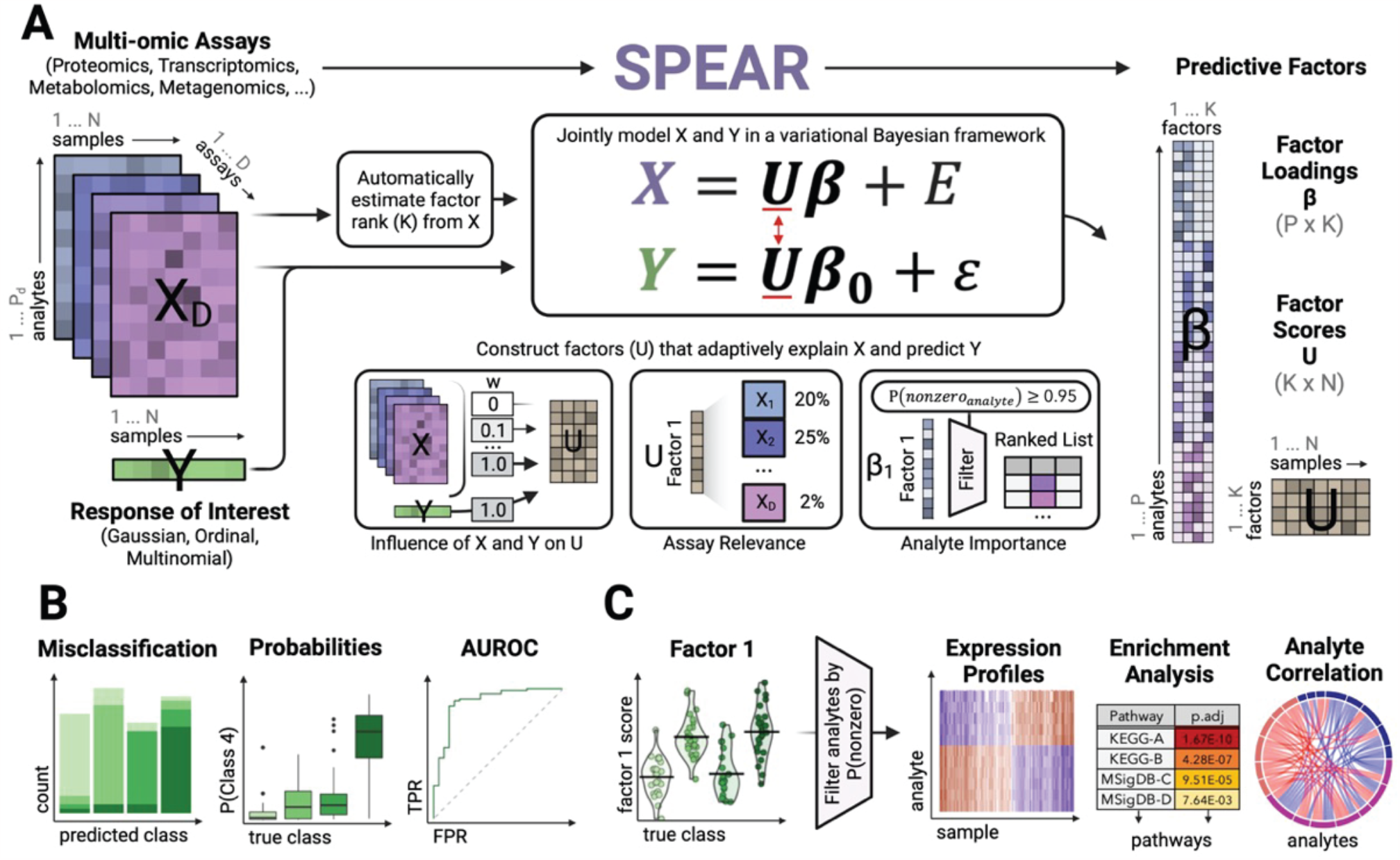
SPEAR workflow overview: (A) SPEAR takes multi-omics data (X) taken from the same N samples, as well as a response of interest (Y). SPEAR supports Gaussian, ordinal, and multinomial types of responses. From these inputs, the algorithm first automatically estimates the minimum number of factors to use in the SPEAR model. X and Y are then jointly modeled in a variational Bayesian framework to adaptively construct factor loadings (β) and scores (U) that explain variance of X and are predictive of Y (reflected in β_0). (B) SPEAR factors are used to predict Y and provide probabilities of class assignment for ordinal and multinomial responses. (C) Downstream biological interpretation of factors is facilitated via automatic feature selection, expression profile analysis, enrichment analysis, and analyte correlation.

Let *X* be an *N* × *p* matrix representing the concatenation of multiple multi-omics assays for *N* samples and *p* features and let *Y* be a length *N* vector representing the response of interest. We assume that *X* is driven by underlying low dimensional factors via linear modeling, coupled with unstructured Gaussian noise (*E*):

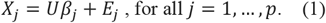

The goal of latent factor model analysis is to decompose *X* into latent factors (*U*) and factor loadings (*β*_*j*_ for all j = 1, …, p). This has been previously accomplished by finding the Bayesian posteriors of *U* and *β* in the probability model (Argelaguet *et al*., 2020) but has not yet been extended to work with a response of interest. Let *Y* be a length *N* vector representing a univariate Gaussian response of interest (see Supplementary Methods C for extensions to ordinal, multinomial, binomial, and multiple responses). Like *X*, let *Y* also be constructed by factors via linear modeling:

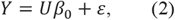

where *β*_0_ is a vector of response coefficients used to construct *Y*. Factors that are most influential for the prediction of *Y* are indicated by larger magnitudes in corresponding *β*_0_.

SPEAR prioritizes the estimation of predictive factors by jointly modeling *X* and *Y* and considering a weighted probability model where the weight parameter (*w*) indicates the emphasis on exploring existing factor structure in *X*:

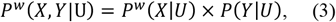

where P^*w*^ (*X*|*U*) is a weighted probabilistic distribution of *X* given *U* and P(*Y*|*U*) is the probabilistic distribution of Y given U. When *w* is large (*w*≥ 1), SPEAR will emphasize the construction of factors that explain the structure of the multi-omics assays (*X*). Intermediary weights (1 > *w* > 0) correspond to a gradual shift in emphasis of predicting the response (*Y*) over explaining assay variance. When *w* is smallest (*w* ≈ 0), SPEAR will forgo any attempt to reconstruct factors from the data and focus entirely on optimizing the prediction of the response. SPEAR automatically selects the weight *w* via cross-validation to balance both a high-prediction accuracy and explainable factors (when these are supported by the data) (Materials and Methods; Supplementary Methods A2).

## 3 Results

### 3.1 SPEAR improves prediction of synthetic responses from simulated multi-omics data

We first evaluated the ability of SPEAR to predict a Gaussian response on simulated data. In the simulation, five predetermined factor signals (*U*) were used to construct four multi-omics assays (*X*), each with 500 analytes (for a total of 2,000 simulated features), for 500 training and 2,000 test samples. The first two simulated factors were assigned to be the multi-omics signatures and were used to construct a Gaussian response vector (*Y*) for both the training and test groups, whereas the remaining three simulated factors were only used to construct *X*. This procedure was repeated across a gradient of signal-to-noise ratios (low signal, moderate signal, and high signal), with 10 independent iterations for each ratio (Supplementary Methods D).

We trained SPEAR across a gradient of values for weight *w* to demonstrate the effect of *w* on SPEAR’s performance. As a comparison, we also employed a two-step MOFA-based model that performs Lasso regression (Tibshirani, 1996) using factors derived from MOFA (denoted as MOFA in the following) as well as a vanilla Lasso regression model using concatenated features in lieu of factors (denoted as Lasso). Model comparison was measured by calculating the mean squared error (MSE) of the test data as well as Pearson correlation testing of the constructed factors against both *Y* (Fig. 2b) and *U* (Fig. 2c).

**Figure 2.**
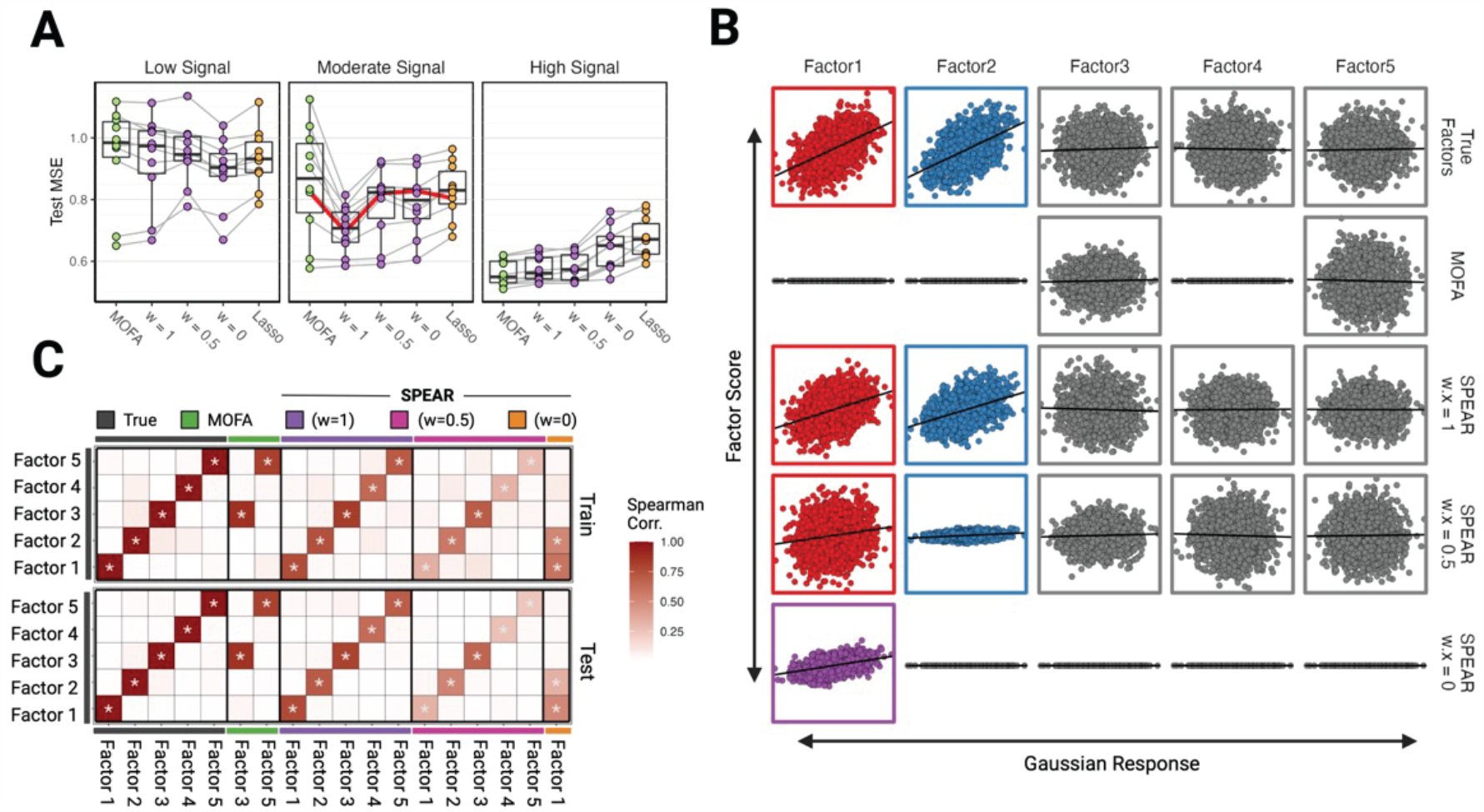
Gaussian simulation results. (A) Boxplots of mean-squared errors of the models on the testing data. MSE results for each simulated iteration are connected. Results are shown for varying signal-to-noise ratios, including low, moderate, and high signals. (B) Scatterplots of various factor scores (y-axis) against the true Gaussian response (x-axis) of the moderate signal test data. Color is applied to factor scores found to be correlated with true factors 1 (red) and 2 (blue) with true factors 3-5 designated as grey. (C) Correlation matrix showing the Spearman correlation between each derived factor of the true factors for both the training and testing data of the moderate signal simulated dataset. Significant correlations are denoted with ‘*’ for p ≤ 0.001.

In scenarios with moderate signal-to-noise, SPEAR with *w* = 1 significantly outperformed MOFA (Fig. 2a). Correlation analyses of the factor scores for one iteration (denoted by the red line in Figure 2a) confirmed that SPEAR with *w* = 1 correctly identified all five simulated factors, whereas MOFA only identified two of the uncorrelated factors (Factor 3 and Factor 5) likely due to the lack of supervision (Fig. 2c), which was confirmed across all 10 iterations (Supplementary Fig. 2). As expected, SPEAR with *w* = 0 condensed all predictive signals into a single factor, as evidenced by its correlation with the first two simulated factors. Overall, SPEAR with higher weights achieved the best predictive performance due to better reconstruction of the multiple underlying simulated factors.

Finally, we repeated the above protocol to test the ability of SPEAR for predicting various non-Gaussian responses (Supplementary Methods A1). Simulated factor construction followed the same protocol above, with only the first two of five factors containing nonlinear signals that were predictive of the response (Supplementary Fig. 3A-B, Supplementary Fig. 4A-B). When the response type was modeled properly (e.g. Gaussian, multinomial, ordinal), SPEAR achieved the best performance via balanced misclassification errors (Supplementary Fig. 3C-D, Supplementary Fig. 4C-D).

**Figure 3.**
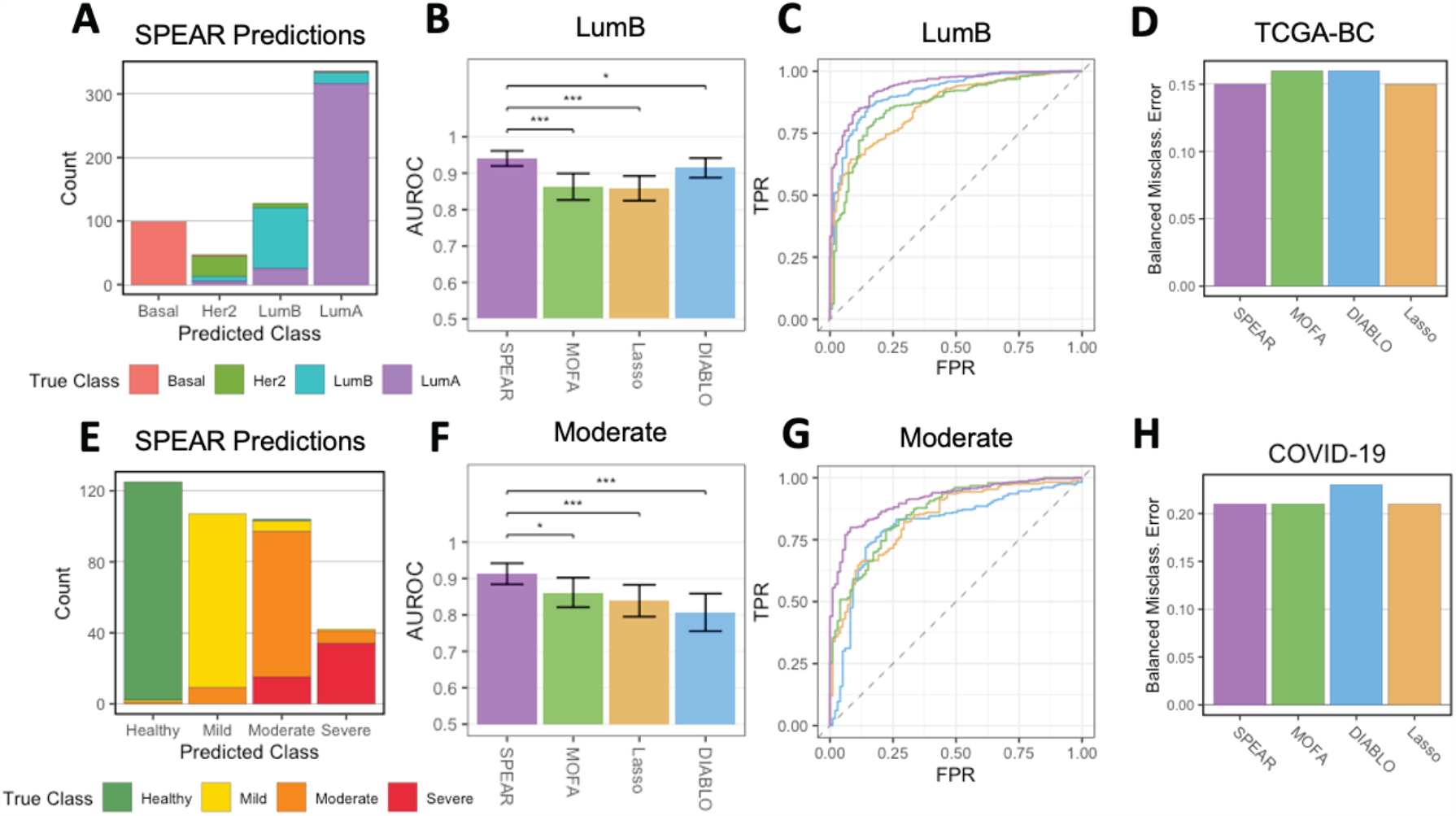
TCGA-BC Tumor Subtype and COVID-19 Severity Prediction Results. (A, E) Test sample class predictions of the SPEAR model, colored by true class. (B, F) Multi-class AUROC statistics for each model for the LumB and Moderate classes. Error bars show the 95% confidence interval found via 2,000 stratified bootstrapping replicates. Significance testing is denoted as * (p ≤ .05), ** (p ≤ .005), and *** (p ≤ .0005). (C) AUROC plot for all models predicting LumB subtype. (G) AUROC plot for all models predicting the moderate severity class. (D, H) Balanced misclassification errors of SPEAR, MOFA, DIABLO, and Lasso on test samples from the (D) TCGA (Breast Cancer) dataset and (H) COVID-19 dataset.

**Figure 4.**
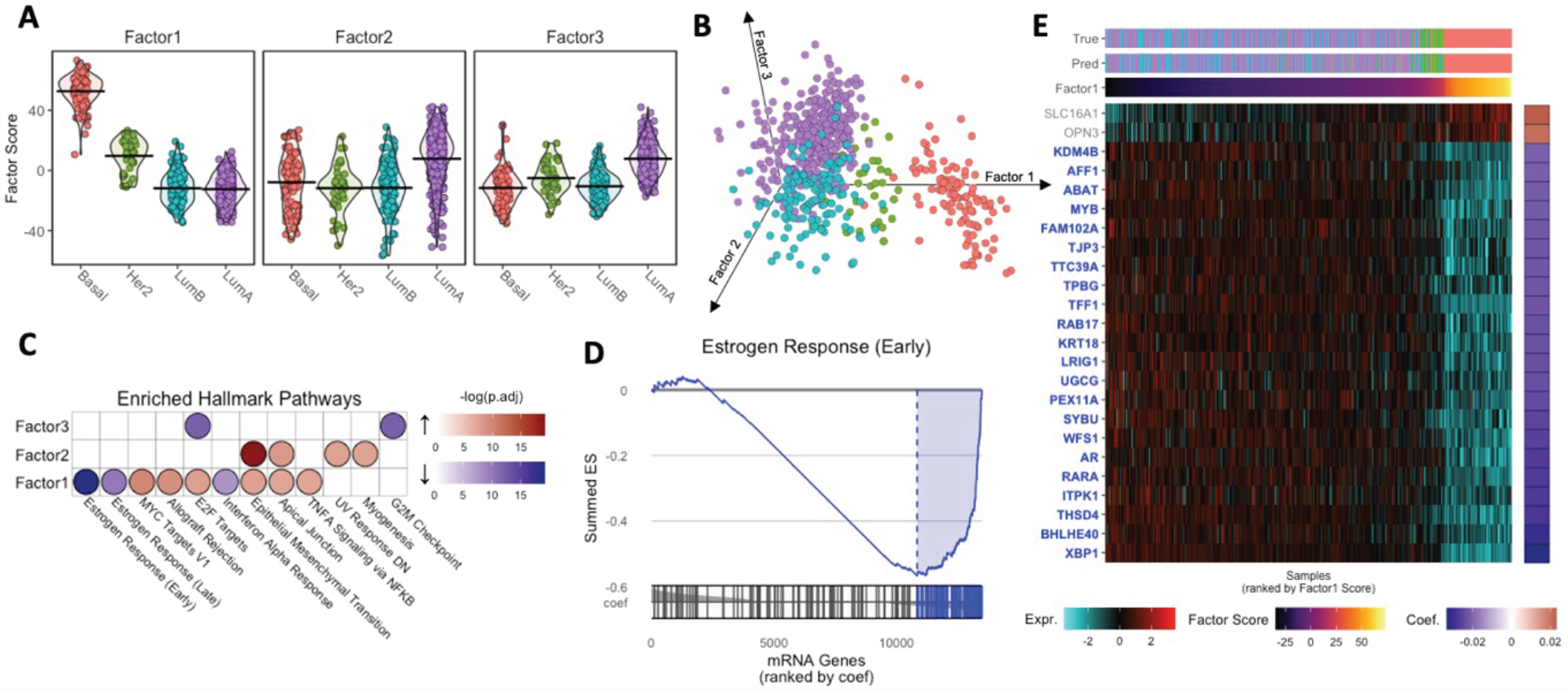
Downstream TCGA-BC Analysis. (A) Grouped violin plot of factors 1-3 scores (y-axis) and tumor subtype (x-axis), with group means marked with a line. (B) 3D scatter plot, embedding samples by factors 1, 2, and 3 scores. Samples are colored by tumor subtype. (C) Dotplot of GSEA results on mRNA features for factors 1-3. Points are shaded by -log(p.adjusted) with color representing enrichment direction. (D) GSEA plot for Estrogen Response (Early) Hallmark pathway for SPEAR Factor 1. mRNA genes are ranked by their assigned projection coefficient from SPEAR factor 1. (E) Heatmap showing normalized expressions for the top 24 mRNA genes involved in the Estrogen Response (Early) Hallmark pathway. mRNA genes were selected with a factor loading (projection coefficient) magnitude ≥ .02. Samples were ranked by Factor 1 score (x-axis) and genes were ranked by projection coefficient (y-axis). Also shown are corresponding true tumor subtypes (True) and SPEAR-predicted tumor subtypes (Pred).

### 3.2 SPEAR improves prediction of breast cancer tumor subtypes and COVID-19 severity

To test whether SPEAR could achieve competitive performance in real data, we applied SPEAR to two publicly available multi-omics datasets: a breast cancer dataset of tumor samples by *Singh et al*. (Singh *et al*., 2019) with the goal of tumor subtype prediction, and a SARS-CoV-2 patient dataset of blood samples by *Yapeng et al*. (Su *et al*., 2020) with the goal of predicting disease severity. We refer to these datasets as TCGA-BC (The Cancer Genome Atlas (Cancer Genome Atlas Network, 2012) - Breast Cancer) and COVID-19, respectively. The TCGA-BC dataset is composed of 989 biopsy samples each associated with RNA-Seq, miRNA, and methylation probe data from primary solid breast cancer tumors that have been annotated according to the PAM50 subtype signature into one of four subtypes: Luminal A (LumA), Luminal B (LumB), HER2-enriched (HER2) and Basal-like (Basal) (Parker *et al*., 2009). The COVID-19 dataset contains plasma protein and metabolite compositions for 254 SARS-CoV-2 positive patient samples and 124 matched healthy subject samples. Each subject is associated with a severity score based on the World Health Organization (WHO) ordinal severity score (Rubio-Rivas *et al*., 2022), binned into four ordinal classes (healthy, mild, moderate, and severe).

We applied SPEAR, Lasso, MOFA, and DIABLO to predict the response from each multi-omics dataset. Datasets were preprocessed and split into training and testing cohorts (see Materials and Methods). Classification performance was evaluated via the balanced misclassification error rate and the area under the receiver operating characteristic (AUROC) curves. AUROC significance was calculated via a bootstrapping procedure and compared via DeLong’s test (DeLong *et al*., 1988).

The advantage of SPEAR was clear when looking at the multi-class AUROC, measuring a classifier’s ability to discriminate each class individually across a gradient of threshold values. SPEAR showed higher AUROC in discriminating between all classes and was significantly better for multiple classes: SPEAR was signifiantly better at predicting both the LumB subtypes from the TCGA-BC dataset (Fig. 3b, Fig. 3c), and moderate SARS-CoV-2 severity from the COVID-19 dataset (Fig. 3f, Fig. 3g) compared to all other models via AUROC comparison testing (LumB – MOFA: p.adj=7.8e-09, Lasso: p.adj=2.1e-07, DIABLO: p.adj=1.9e-02; Moderate – MOFA: p.adj=1.4e-02, Lasso: p.adj=1.9e-04, DIABLO: p.adj=1.5e-05). Upon investigation, the improved predictive performance of SPEAR on these classes was not due to a single factor but was rather achieved through combining information from multiple multi-omics factors (Supplementary Fig. 5, Supplementary Fig. 6). The balanced misclassification errors of SPEAR (0.15, 0.21) were comparable to those of Lasso (0.15, 0.21), MOFA (0.16, 0.21), and DIABLO (0.16, 0.23) for the TCGA-BC dataset and COVID-19 dataset respectively (Fig. 3a, Fig. 3e, Fig. 3d, Fig. 3h). Overall, our results demonstrate that SPEAR outperforms current state-of-the-art methods for predicting a response using multi-omics data.

**Figure 5.**
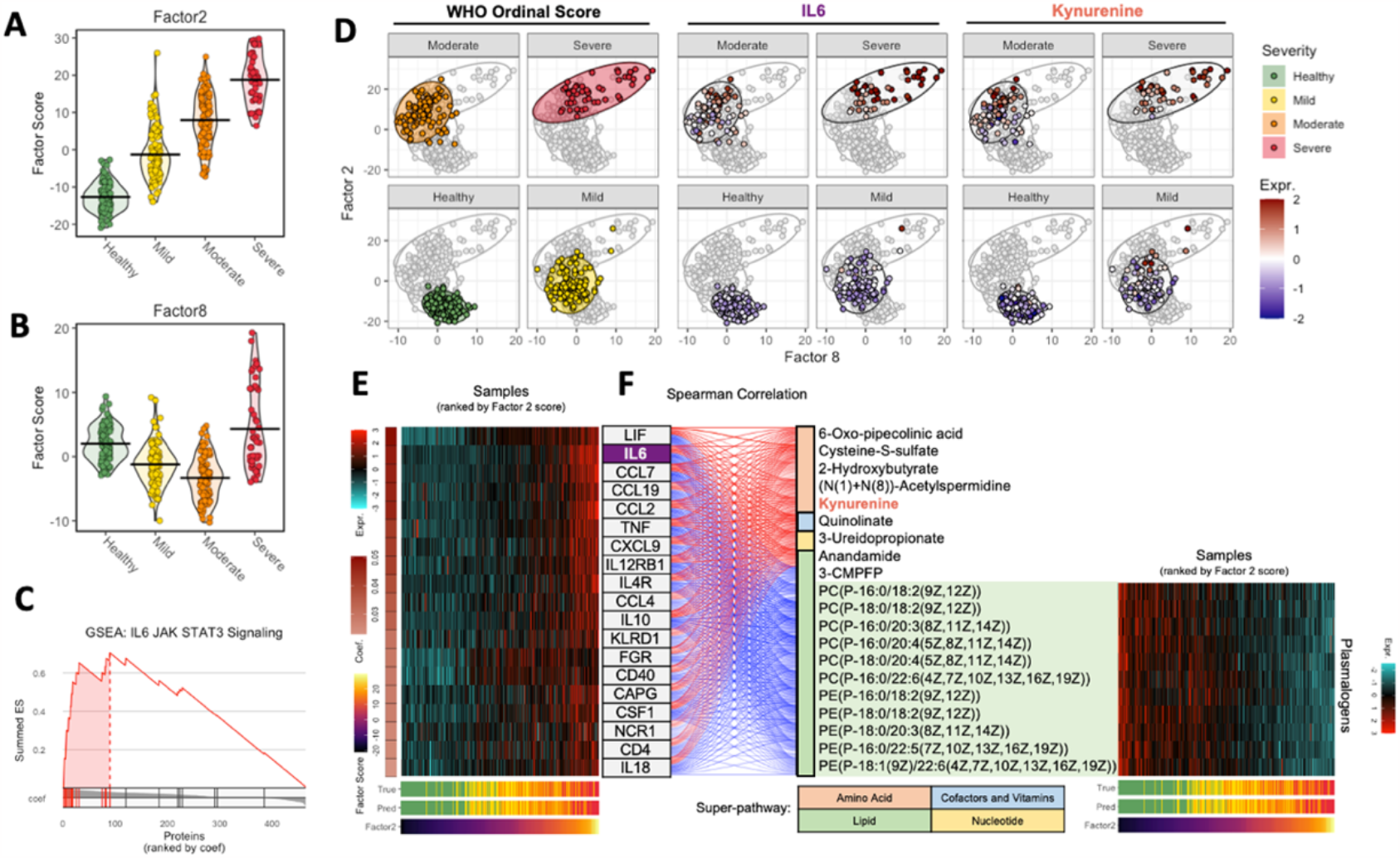
Downstream COVID-19 Dataset Analysis. (A) Grouped violin plot of factor 2 scores (y-axis) and simplified WHO score (x-axis), with group means marked with a line. (B) Grouped violin plot of factor 8 scores (y-axis) and simplified WHO score (x-axis), with group means marked with a line. (C) GSEA plot for IL6 JAK STAT3 Signaling Hallmark pathway for SPEAR Factor 2. Proteins are ranked by their assigned projection coefficient from SPEAR factor 2. (D) Embedding of samples by factor 8 (x-axis) and factor 2 (y-axis) scores. Samples are colored by WHO Ordinal Score, normalized IL6 expression, and normalized kynurenine expression. (E) Heatmap showing normalized expressions for proteins involved in the IL6 JAK STAT3 Signaling Hallmark pathway. Samples were ranked by Factor 2 score (x-axis) and proteins were ranked by projection coefficient (y-axis). Also shown are corresponding true patient severity scores (True) and SPEAR (sd) predicted severity scores (Pred). (F) Alluvial plot showing correlation between the IL6 JAK STAT3 proteins and the top metabolite features contributing to SPEAR Factor 2. Metabolites are grouped by positive/negative correlation and super-pathway. Also shown are normalized plasmalogen expressions of samples ranked by Factor 2 score (x-axis).

It was notable that the balanced misclassification error of SPEAR was comparable with other methods even though SPEAR does not optimize purely for predictive performance. Rather, SPEAR favors multi-omics assay influence in the construction of predictive factors by choosing the largest *w* whose mean cross-validated error falls within one standard deviation of the overall minimum cross-validated error. If we instead choose the *w* that minimizes the cross-validated error (denoted as SPEAR *min*), the balanced misclassification error is better than all other models on both the TCGA-BC and COVID-19 datasets (0.13, 0.19) (Supplementary Fig. 7a, 7d). Similarly, this approach achieves the best AUROC values for all classes (Supplementary Fig. 7c, 7f). The SPEAR min model showed only a slight improvement in classification accuracy but chose considerably lower w values (w = 0.5 for TCGA-BC and w = 0.01 for COVID-19) than the default SPEAR model (denoted as SPEAR sd when needed for clarity) (w = 2.0 for both datasets) (see Supplementary Fig. 8). Although we recommend and used the default SPEAR model to enhance the downstream interpretation, SPEAR *min* can be used in cases where prediction is the key objective regardless of multi-omics assay influence in factor construction.

### 3.3 SPEAR identifies steroid response pathways associated with TCGA PAM50 subtypes

Several factors returned by SPEAR were associated with different biological pathways that distinguish the PAM50 subtypes. Factors 1-3 clearly distinguished one or more subtypes, which motivated us to investigate what biological pathways each factor was associated with (Fig. 4, Supplementary Fig. 4). Factor 1 was strongly associated with the Basal subtype and moderately associated with the HER2e subtype (Fig. 4a, Fig. 4b), while the negative factor loadings were enriched for Estrogen Response (ER) (Early and Late) pathways of the Molecular Signatures Database (MSigDB) Hallmark collection (Liberzon *et al*., 2015) (Fig. 4c, Fig. 4d).

The anti-correlation between the ER pathways and the Basal subtype, the highest scoring Factor 1 subtype, reflects that it is most strongly associated with a triple-negative profile for Estrogen, Progesterone, and the HER2e receptor (Prat *et al*., 2013). Whereas HER2e-classified samples were predominantly hormone receptor (HR) negative, a proportion is also HR positive (Bastien *et al*., 2012; Prat *et al*., 2015). LumA and LumB are hormone receptor positive (HR) and retain expression of ER, and the Progesterone receptor (PR) in the case of LumA, and in some proportion of LumB (Prat *et al*., 2015). Indeed, genes *XBP1, AGR2*, and *CA12* which yielded high posterior selection probabilities as well as the strongest negative projection coefficients in Factor 1 (Fig. 4e, Supplemental Table 1) for the Estrogen Response (Late) pathway were all associated with a breast cancer Steroid Responsiveness (SR) module that indicates functional steroid response (Fredlund *et al*., 2012). Genes *XBP1* and *MYB* were in the top 20% of the ER Early and Late pathway genes with respect to the SPEAR projection coefficient magnitude. *MYB* is a direct target of Estrogen signaling and is overexpressed in most ER+ cancers (Gonda *et al*., 2008), and both genes were also identified as part of a luminal expression signature (Cancer Genome Atlas Network, 2012).

The positive association of Factor 1 with the MYC Targets V1 Hallmark pathway is consistent with the association of MYC signaling and Basal subtypes identified in earlier TCGA and other analyses (Xu *et al*., 2010). Additionally, four of the miRNAs associated with Factor 1, *hsa-mir-18a, hsamir-10a, hsa-mir135b, hsa-mir-577*, were identified in a miRNA analysis of TCGA data to have diagnostic significance for triple-negative breast cancer (Fan and Liu, 2019). *Hsa-mir-18a* has been shown in independent datasets to downregulate ERα (Klinge, 2012) and is associated with worse overall survival (Luengo-Gil *et al*., 2019). Hypomethylation of *MIA*, a PAM50 gene, which has the second highest in magnitude SPEAR projection coefficient for Factor 1, has been associated with the Basal subtype in TCGA and independent data (Bardowell *et al*., 2013).

Factor 1 distinguished Basal-like and HER2e subtypes from LumA/LumB, while Factors 2 and 3 distinguished LumA from the other subtypes. Pathways enriched in Factor 2 were associated with genes that are more highly expressed in LumA, whereas pathways in Factor 3 were associated with genes that are downregulated in LumA (Fig. 4a, Fig. 4b). The strongest signal in Factor 2 was in the Epithelial Mesenchymal Transition (EMT) Hallmark pathway, and in Factor 3 the G2M Checkpoint Hallmark pathway (Fig. 4c). It has been previously observed that LumB tumors are enriched in proliferation and cell-cycle associated genes in comparison to LumA (Prat *et al*., 2015), whereas the positive association of LumA with an EMT phenotype is somewhat surprising, as LumA tumors are generally considered to be associated with a stronger epithelial phenotype in comparison to the Basal-like subtype (Felipe Lima *et al*., 2016). The role of EMT has been primarily studied in non-luminal tumors^28^, and therefore the presence of this pathway distinguishing the subtypes warrants further investigation with respect to the underlying biology. Interestingly, the miRNAs with the top magnitude projection coefficients in Factor 2 and which have a positive association with the factor, *hsa-mir199a/b*, have been associated with EMT (Drago-García *et al*., 2017; Wang *et al*., 2019). Overall, the molecular signatures identified via SPEAR factorization of the TCGA-BC data provided both well-documented and novel associations with PAM50 breast cancer subtypes.

### 3.4 SPEAR identifies multi-omics factors and pathways associated with COVID-19 severity

On the COVID-19 dataset, SPEAR identified several factors that were significantly associated with the WHO ordinal severity score (Fig. 5, Supplementary Fig. 5). Investigation of the association of each factor with the COVID-19 severity revealed that the Factor 2 score showed a positive ordinal association (Fig. 5a). SPEAR also identified several factors that were heavily associated with identifying COVID-19 severities, including Factor 1 score for mild severity and Factor 8 for moderate severity (Fig. 5b, Supplementary Fig. 4). Embedding the samples by Factors 2 and 8 scores revealed a trajectory for SARS-CoV-2 severities (Fig. 5d). Calculation of the variance explained for these three factors revealed that Factor 2 had the largest influence from both assays (16% proteomics, 11% metabolomics) compared to Factor 1 (20% proteomics, 1% metabolomics) and Factor 8 (1% proteomics, 2% metabolomics) (Supplementary Fig. 5). We opted to further investigate Factor 2 due to its larger multi-omics influences.

Proteomic enrichment analysis of Factor 2 identified several significant pathways from the MSigDB Hallmark pathway database (Supplemental Table 2). One pathway enriched in Factor 2 was the Janus kinase Signal Transducer and Activator of Transcription (JAK-STAT) signaling pathway (Seif *et al*., 2017) (Fig. 5c). Pathway member Interleukin 6 (IL-6) was also found to be a key contributor to the Factor 2 score, with the second highest projection coefficient (Fig. 5e, Supplemental Table 3). IL-6 is a proinflammatory cytokine proposed as an inflammatory biomarker for COVID-19 severity (Azevedo *et al*., 2021) and is an activator of the JAK-STAT signaling pathway. The JAK-STAT signaling pathway is involved in immune regulation, lymphocyte growth and differentiation, and promotes oxidative stress, serving as an attractive therapeutic target for COVID-19 treatment (Luo *et al*., 2020).

The Factor 2 metabolic signature, extracted via automatic feature selection of the top contributing metabolites to the Factor 2 score, identified members of the amino acid, cofactors and vitamins, lipid, and nucleotide super-pathways (Fig. 5f). Kynurenine, a key contributor to the Factor 2 score, is a known marker of severe/fatal COVID-19 trajectory (Danlos *et al*., 2021; Mangge *et al*., 2021). The tryptophan/kynurenine (Trp/Kyn) pathway is activated by inflammatory cytokines (Almulla *et al*., 2022), which is consistent with its positive correlation with the cytokines of Factor 2 such as IL-6 (Fig. 5f).

Several plasmalogens, including phosphatidylcholines (PCs) and phosphatidylethanolamines (PEs) were also found to be inversely associated with the ordinal severity trend of Factor 2 (Fig. 5f). Plasmalogens are plasma-borne antioxidant phospholipids that provide endothelial protection during oxidative stress (Messias *et al*., 2018), which could account for the inverse association with the JAK-STAT signaling pathway proteins. Our multi-omics results further support the utility of plasmalogens as prognostic indicators of COVID-19 severity (Pike *et al*., 2022).

## 4 Discussion

SPEAR extracts interpretable multi-omics signatures from predictive factors via supervised dimensionality reduction and can model multiple types of responses, including Gaussian, multinomial, and ordinal. We have compared SPEAR to state-of-art linear dimension reduction methods as well as direct predictive models via Lasso regression in both simulated and real datasets. Unlike similar probabilistic factor models such as MOFA and iClusterBayes, the supervised SPEAR framework integrates the response of interest into the dimensionality reduction, resulting in factors containing signals from both the multi-omics assays and the response when supported by the data. In addition, SPEAR estimates analyte significance probabilities per factor, eliminating the need to perform feature selection via the loading coefficients entirely.

Defining the number of factors to be constructed can be critical when applying factor model-based methods. We have observed that underestimating the rank of a SPEAR model can lead to poor performance at higher values of *w*. To this end, SPEAR begins by estimating the minimum number of factors required to accurately represent a multi-omics dataset. This adaptive factor number estimation approach is not restrictive to SPEAR and can be incorporated into other dimensionality reduction pipelines where the optimal number of factors to use is unknown.

While generalizable to many types of multi-omics, SPEAR does assume a linear dependence of the latent factors on the features. As such, the application of SPEAR to nonlinear analyte dependence may produce undesirable results. This could be mediated by appropriately preprocessing nonnormal multi-omics assays through transformations. A more data-adaptive solution would be to model the non-linear relationship between the features and factors via the generalized additive model (Hastie and Tibshirani, 1995).

In conclusion, the SPEAR model decomposes high-dimensional multi-omics datasets into interpretable low-dimensional factors with high predictive power without the need for parameter tuning. SPEAR returns both sparse (regression) and full (projection) coefficients as well as feature-wise posterior probabilities used to assign analyte significance. SPEAR is currently hosted as a publicly available R-package at https://bitbucket.org/kleinstein/SPEAR.

## 5 Online Methods

### 5.1 Automated selection of weight parameter (w) via cross-validation (CV)

SPEAR controls the level of prioritization in construction of predictive factors by adjusting the importance of the fit towards *X* through the weight parameter *w*. A low *w* (*w* ≈ 0) will yield factors that emphasize prediction of *Y*, while higher values for *w* (*w*≥ 1) will construct factors that also can explain the structure of *X*.

We propose to automatically balance between the construction of factors and the prediction of the response through the weight parameter w, which is selected via cross-validation. SPEAR factors are constructed *K* times, where *K* is the number of folds used to split the training data, and the mean cross-validated error is calculated per weight *w* across folds. By default, the largest *w* whose mean cross-validated error falls within one standard deviation of the overall minimum cross-validated error is selected (Hastie *et al*., 2009), favoring higher values of *w* by design. Thus, SPEAR constructs predictive factors influenced by the multi-omics assays if data supported, returning a simple linear predictive model otherwise.

### 5.2 Automatic Relevance Determination (ARD), spike-and-slab priors and feature selection

Multi-omics datasets are often high-dimensional, and different assays may capture different sources of variations. To combat this, SPEAR models assay-wise factor influence using Automatic Relevance Determination (ARD) (Wipf and Nagarajan, 2007) and the feature sparsity using spike-and-slab priors (Supplementary Methods A1). ARD allows for the construction of factors with asymmetrical assay contributions, and spike-and-slab priors identify key sparse predictive multi-omics features. Like alternative sparse supervised methods, feature importance is represented by a vector of coefficients for each factor, with a higher magnitude representing a higher contribution to the factor construction.

In contrast to methods like MOFA and DIABLO, SPEAR returns two different types of coefficients for each multi-omics feature, denoted as regression coefficients and projection coefficients. The regression coefficients represent the sparse predictive features most important for factor construction, and the projection coefficients capture a factor’s influence on all analytes. These projection coefficients are intended for downstream factor interpretation where seemingly redundant features ignored in the projection coefficients are vital to understanding the underlying biology^12^. Along with the coefficients, SPEAR will also approximate the posterior probabilities of multi-omics features being non-zero, allowing for automatic feature selection by keeping features with a posterior probability of being non-zero above 0.95, which is analogous to a false discovery rate (FDR) of 0.05. Choosing features based on this estimated posterior probability provides an advantage over other factor models, which facilitate feature selection by arbitrary cut-offs on coefficient values.

### 5.3 Selection of SPEAR model rank

The model rank, or number of factors to construct, need to be determined by the user, and proper choices of them are data dependent. Alternative factor models utilize a static number such as 15 or 20 factors as a default. This is problematic in predictive settings, as underestimating the required number of predictive factors will result in worse performance. To this end, SPEAR begins by suggesting a sufficient model rank by estimating how the largest number estimable factors from the multi-omics assays whose influence on the data could not be accounted for by pure noise, based on a data randomization approach used in the regression problem (Guan and Tibshirani, 2020; Tian, 2020). This procedure is explained in detail in Supplementary Methods A2.

### 5.4 TCGA-BC dataset background and preprocessing

The TCGA-BC dataset was adapted from *Singh et al*. (Singh *et al*., 2019), consisting of 16851 mRNAs, 349 miRNAs, and 9482 CpG methylation sites. Each sample was taken from a primary solid breast cancer tumor that has been classified according to the PAM50 subtype signature, a 50-gene signature. The PAM50 signature is one of the main intrinsic breast cancer signatures used in clinical practice to determine the course of therapy (Wallden *et al*., 2015; Goldhirsch *et al*., 2013). *Singh et al*. showed that even when the 50 genes were removed from the dataset, it was still possible to predict subtype class using the remaining multi-omics features, indicating that there are underlying biological pathways that can distinguish and drive the subtypes.

While most preprocessing was kept as described in *Singh et al*., including the removal of the PAM50 signature mRNA genes, we additionally removed the least 20% variable features from each assay to reduce the total number of features. We did not exclude the PAM50 genes from the methylation results as the markers are used in gene expression assays. Samples were then split into train (*n*_*train*_ = 379) and test (*n*_*test*_ = 610) sets as described in *Singh et al*. Finally, all three assays were scaled and centered to be normally distributed (μ = 0, σ^2^ = 1).

### 5.4 COVID-19 dataset background and preprocessing

The SARS-CoV-2 (COVID-19) dataset contains multi-omics data from 254 samples from SARS-CoV-2 positive patients and 124 samples from matched healthy subjects from *Yapeng et. al*. (Su *et al*., 2020). Samples were taken from one of two timepoints, T1 and T2. At each timepoint, participants were assigned a COVID-19 severity score based on a World Health Organization (WHO) ordinal scale for clinical improvement: uninfected (0), ambulatory without (1) and with activity limitation (2), hospitalized without (3) and with oxygen therapy (4), hospitalized with non-invasive ventilation (5), intubation (6) or ventilation with additional organ support (7) and death (8). These ordinal values were condensed into four classes: healthy (0), mild (1-2), moderate (3-4), and severe (5-7).

MissForest, a non-parametric missing value imputation for mixed-type data was utilized to address missing values in both the proteomics and metabolomics (Stekhoven and Bühlmann, 2012). The metabolite expression values required quantile normalization and log transformation. Both proteomic and metabolomic expression values were then scaled and centered to be normally distributed (μ = 0, σ^2^ = 1).

## Acknowledgements

We are grateful to Joann Diray-Arce for her assistance with the metabolomic enrichment analysis for the COVID-19 dataset. We also acknowledge Amrit Singh, Casey Shannon, and Scott Tebbutt for the helpful conversations and support with DIABLO.

## References

Almulla, A.F. et al. (2022) The tryptophan catabolite or kynurenine pathway in COVID-19 and critical COVID-19: a systematic review and meta-analysis. BMC Infect Dis, 22, 615.

Argelaguet, R. et al. (2020) MOFA+: a statistical framework for comprehensive integration of multi-modal single-cell data. Genome Biol, 21, 111.

Azevedo, R.B. et al. (2021) Covid-19 and the cardiovascular system: a comprehensive review. J Hum Hypertens, 35, 4–11.

Banoth, B. and Cassel, S.L. (2018) Mitochondria in innate immune signaling. Transl Res, 202, 52–68.

Bardowell, S.A. et al. (2013) Differential methylation relative to breast cancer subtype and matched normal tissue reveals distinct patterns. Breast Cancer Res Treat, 142, 365–380.

Bastien, R.R.L. et al. (2012) PAM50 breast cancer subtyping by RT-qPCR and concordance with standard clinical molecular markers. BMC Med Genomics, 5, 44.

Bhattacharya, S. et al. (2014) ImmPort: disseminating data to the public for the future of immunology. Immunol Res, 58, 234–239.

Bodkin, N. et al. (2022) Systematic comparison of published host gene expression signatures for bacterial/viral discrimination. Genome Med, 14, 18.

Bolen, C.R. et al. (2014) Dynamic expression profiling of type I and type III interferon-stimulated hepatocytes reveals a stable hierarchy of gene expression. Hepatology, 59, 1262–1272.

Cancer Genome Atlas Network (2012) Comprehensive molecular portraits of human breast tumours. Nature, 490, 61–70.

Cantini, L. et al. (2021) Benchmarking joint multi-omics dimensionality reduction approaches for the study of cancer. Nat Commun, 12, 124.

Chawla, D.G. et al. (2022) Benchmarking transcriptional host response signatures for infection diagnosis. Cell Syst, 13, 974–988.e7.

Danlos, F.-X. et al. (2021) Metabolomic analyses of COVID-19 patients unravel stage-dependent and prognostic biomarkers. Cell Death Dis, 12, 1–11.

DeLong, E.R. et al. (1988) Comparing the Areas under Two or More Correlated Receiver Operating Characteristic Curves: A Nonparametric Approach. Biometrics, 44, 837–845.

Drago-García, D. et al. (2017) Network analysis of EMT and MET micro-RNA regulation in breast cancer. Sci Rep, 7, 13534.

Fan, C. and Liu, N. (2019) Identification of dysregulated microRNAs associated with diagnosis and prognosis in triple-negative breast cancer: An in silico study. Oncol Rep, 41, 3313–3324.

Felipe Lima, J. et al. (2016) EMT in Breast Carcinoma-A Review. J Clin Med, 5, E65.

Fourati, S. et al. (2022) Pan-vaccine analysis reveals innate immune endotypes predictive of antibody responses to vaccination. Nature Immunology, 1–11.

Fredlund, E. et al. (2012) The gene expression landscape of breast cancer is shaped by tumor protein p53 status and epithelial-mesenchymal transition. Breast Cancer Res, 14, R113.

Goldhirsch, A. et al. (2013) Personalizing the treatment of women with early breast cancer: highlights of the St Gallen International Expert Consensus on the Primary Therapy of Early Breast Cancer 2013. Ann Oncol, 24, 2206–2223.

Gonda, T.J. et al. (2008) Estrogen and MYB in breast cancer: potential for new therapies. Expert Opin Biol Ther, 8, 713–717.

Guan, L. and Tibshirani, R. (2020) Post model-fitting exploration via a “Next-Door” analysis. Canadian Journal of Statistics, 48, 447–470.

Hagan, T. et al. (2022) Transcriptional atlas of the human immune response to 13 vaccines reveals a common predictor of vaccine-induced antibody responses. Nat Immunol, 23, 1788–1798.

Hastie, T. et al. (2009) The elements of statistical learning: data mining, inference, and prediction Springer.

Hastie, T. and Tibshirani, R. (1995) Generalized additive models for medical research. Stat Methods Med Res, 4, 187–196.

Klinge, C.M. (2012) miRNAs and estrogen action. Trends Endocrinol Metab, 23, 223–233.

Lazear, H.M. et al. (2019) Shared and Distinct Functions of Type I and Type III Interferons. Immunity, 50, 907–923.

Li, W. et al. (2012) Identifying multi-layer gene regulatory modules from multi-dimensional genomic data. Bioinformatics, 28, 2458–2466.

Liberzon, A. et al. (2015) The Molecular Signatures Database (MSigDB) hallmark gene set collection. Cell Syst, 1, 417–425.

Luengo-Gil, G. et al. (2019) Clinical and biological impact of miR-18a expression in breast cancer after neoadjuvant chemotherapy. Cell Oncol (Dordr), 42, 627–644.

Luo, W. et al. (2020) Targeting JAK-STAT Signaling to Control Cytokine Release Syndrome in COVID-19. Trends in Pharmacological Sciences, 41, 531–543.

Mangge, H. et al. (2021) Increased Kynurenine Indicates a Fatal Course of COVID-19. Antioxidants, 10, 1960.

Messias, M.C.F. et al. (2018) Plasmalogen lipids: functional mechanism and their involvement in gastrointestinal cancer. Lipids in Health and Disease, 17, 41.

Mo, Q. et al. (2018) A fully Bayesian latent variable model for integrative clustering analysis of multi-type omics data. Biostatistics, 19, 71–86.

Nakaya, H.I. et al. (2011) Systems biology of vaccination for seasonal influenza in humans. Nat Immunol, 12, 786–795.

Parker, J.S. et al. (2009) Supervised risk predictor of breast cancer based on intrinsic subtypes. J Clin Oncol, 27, 1160–1167.

Pike, D.P. et al. (2022) Plasmalogen Loss in Sepsis and SARS-CoV-2 Infection. Frontiers in Cell and Developmental Biology, 10.

Prat, A. et al. (2015) Clinical implications of the intrinsic molecular subtypes of breast cancer. Breast, 24 Suppl 2, S26–35.

Prat, A. et al. (2013) Molecular characterization of basal-like and non-basal-like triple-negative breast cancer. Oncologist, 18, 123–133.

Ramilo, O. et al. (2007) Gene expression patterns in blood leukocytes discriminate patients with acute infections. Blood, 109, 2066–2077.

Rubio-Rivas, M. et al. (2022) WHO Ordinal Scale and Inflammation Risk Categories in COVID-19. Comparative Study of the Severity Scales. J Gen Intern Med, 37, 1980–1987.

Seif, F. et al. (2017) The role of JAK-STAT signaling pathway and its regulators in the fate of T helper cells. Cell Communication and Signaling, 15, 23.

Singh, A. et al. (2019) DIABLO: an integrative approach for identifying key molecular drivers from multi-omics assays. Bioinformatics, 35, 3055–3062.

Stekhoven, D.J. and Bühlmann, P. (2012) MissForest—non-parametric missing value imputation for mixed-type data. Bioinformatics, 28, 112–118.

Su, Y. et al. (2020) Multi-Omics Resolves a Sharp Disease-State Shift between Mild and Moderate COVID-19. Cell, 183, 1479–1495.e20.

Tenenhaus, A. et al. (2014) Variable selection for generalized canonical correlation analysis. Biostatistics, 15, 569–583.

Tenenhaus, A. and Tenenhaus, M. (2011) Regularized Generalized Canonical Correlation Analysis. Psychometrika, 76, 257–284.

Tian, X. (2020) Prediction error after model search. The Annals of Statistics, 48, 763–784.

Tibshirani, R. (1996) Regression shrinkage and selection via the lasso. Journal of the Royal Statistical Society: Series B (Methodological), 58, 267–288.

Walker, K.A. et al. (2023) Proteomics analysis of plasma from middle-aged adults identifies protein markers of dementia risk in later life. Sci Transl Med, 15, eadf5681.

Wallden, B. et al. (2015) Development and verification of the PAM50-based Prosigna breast cancer gene signature assay. BMC Medical Genomics, 8, 54.

Wang, Q. et al. (2019) Overview of microRNA-199a Regulation in Cancer. Cancer Manag Res, 11, 10327–10335.

Wipf, D. and Nagarajan, S. (2007) A New View of Automatic Relevance Determination. In, Advances in Neural Information Processing Systems. Curran Associates, Inc.

Xu, J. et al. (2010) MYC and Breast Cancer. Genes Cancer, 1, 629–640.

Yu, F. et al. (2019) Breast cancer prognosis signature: linking risk stratification to disease subtypes. Brief Bioinform, 20, 2130–2140.

