## Supplementary Methods for "A supervised Bayesian factor model for the identification of multi-omics signatures"

In this supplementary material, we cover (1) full SPEAR model details, (2) algorithm details for variational Bayesian model, and (3) additional empirical results.

**F References** **20**

#### A. SPEAR overview

Fixing the model rank $K$ and the weight parameter $w$, SPEAR considers the following weighted distribution for the data given the latent factor $\mathbf{U}$ and the model parameters:

$$P(\mathbf{X}, \mathbf{Y}|\mathbf{U}, \Theta) = \left(\prod_{i=1}^{N}\prod_{j=1}^{p} P^{w}(x_{i,j}|\mathbf{u}_i, \Theta)\right) \times \left(\prod_{i=1}^{N}\prod_{j=1}^{p_0} P(y_{i,j}|\mathbf{u}_i, \Theta)\right). \tag{1}$$

For the latent factor $\mathbf{U}$, we model it as a noisy realization of some linear function of $\mathbf{Z}$, and let

$$\mathbf{U} = \mathbf{Z}\boldsymbol{\beta} + \boldsymbol{\mathcal{E}}, \tag{2}$$

where $\boldsymbol{\mathcal{E}}_{ik} \sim \mathcal{N}(0, 1)$. We introduce $\boldsymbol{\mathcal{E}}$ here for stability that can happen when $Z\boldsymbol{\beta}$ becomes very small. $\Theta = \left\{\sigma^2, \boldsymbol{\beta}, \mathbf{B}\right\}$ is the collection of all parameters. Let $\Gamma(.|a, c)$ and $\text{Beta}(.|a, c)$ represent the Gamma and Beta distributions with parameters $(a, c)$:

$$\begin{aligned}\Gamma(x|a, c) =& \frac{c^{\alpha}}{\Gamma(\alpha)} x^{\alpha-1}\exp(-cx)\\ \text{Beta}(x|a, c) =& \frac{\Gamma(a+c)x^{a-1}(1-x)^{c-1}}{\Gamma(a)\Gamma(c)}\end{aligned}$$

We use the following priors for the model parameters:

- Inverse Gamma prior for variances. Let $\sigma_j^2$ be the variance of $X_j$. For convenience, we let $\nu_j = \frac{1}{\sigma_j^2}$ and will use $\nu_j$ from here.

$$P(\nu_j) = \Gamma(\nu_j|a_0, c_0) \tag{3}$$

- For $j = 1, \ldots, p$, let $\beta_{jk} = \hat{\beta}_{jk}\gamma_{jk}$, where $\hat{\beta}_{jk} \sim \mathcal{N}(0, \frac{1}{\tau_{g_j k}})$ and $\gamma_{jk} \sim Binary(\pi_{g_j k})$. Here $g_j$ is the group id of feature $j$ and we model different groups to have different prior non-zero probability and variance when being non-zero. For example, by default, we set features from the same assay as one group. Symmetrically, we let $B_{jk} = \hat{B}_{jk}s_{jk}$ where $\hat{B}_{jk} \sim \mathcal{N}(0, \frac{1}{\tau_{g_j k}})$ and $s_{jk} \sim Binary(\pi_{g_j k})$ for $j = 1, \ldots, p$.

- We do not impose sparsity for $\overline{B}_{jk}$, $j > p$, which corresponds to projection coefficients of $Y_j$ onto the factors, and model $\overline{B}_{jk} \sim \mathcal{N}(0, \frac{1}{\tau_0})$.

To avoid the need of extensive parameter tuning, we further model the priors $\tau$, $\pi$ with some hyper-prior distributions for group $g = 1, \ldots, G$ and factor $k = 1, \ldots, K$:

- We model $\tau_{gk}$ with a Gamma prior,

$$P(\tau_{gk}) = \Gamma(\tau_{gk}|a_1, c_1).$$

- We model $\pi_{gk}$ with a Beta prior,

$$P(\pi_{gk}) = \text{Beta}(\pi_{gk}|a_2, c_2).$$

The hyper prior modeling requires no user tuning on $\tau_{\ell k}$ or $\pi_{\ell k}$. As a result, such a model is referred to an ARD (automatic relevance determining) model since feature sparsity and effect size for each group is estimated from the data (Wipf and Nagarajan, 2007).

In this paper, we model the features $X$ as Gaussian but allow $Y$ to be either Gaussian, categorical or ordinal, which are several of the most common data types in practice. We can also model $X$ to be different data types, although in practice $X$ is usually continuous or binary. For both cases, modeling $X$ as Gaussian is reasonable for signal extraction purpose. We allow for more flexibility in modeling $Y$ for the purpose of both better signal extraction and interpretability. When $Y$ is a multiclass categorical response or an ordinal response, modeling $Y$ as Gaussian may lead to worse estimation (if $Y$ is highly non-linear in its nominal values in the ordinal case, or having co-linearity in the multinomial case). The predicted value can also be difficult to interpret.

#### A.1. Modeling of various response types

In Eq. (1), $p_0$ is the dimension of response so that we can deal with multi-dimensional response $y$ as well. For convenience, we discuss only the one-dimensional response setting with $p_0 = 1$ and denote $y_{i1}$ by $y_i$. Modeling of the multi-dimensional response is a simple extension of the one-dimensional problem. By taking proper forms for $P(y_i|\mathbf{u}, \Theta)$, we can flexibly model different response types. SPEAR support four response types:

**Continuous response:** We consider a Gaussian model for continuous response $y$:

$$P(y_i|\mathbf{u}_i, \Theta) = \frac{1}{\sqrt{2\pi\bar{\sigma}^2}} \exp(-\frac{(y_i - \mathbf{u}_i^\top \bar{B})^2}{2\bar{\sigma}^2})$$

where $\bar{B}$ is the projection coefficient from $y$ onto $\mathbf{U}$, and $\bar{\sigma}^2$ is the variance of noise in $y$.

**Two-class categorical response** We model the two-class categorical response $y_i \in \{0, 1\}$ using the logistic regression:

$$P(y_i|\mathbf{u}_i, \Theta) = \left[\frac{1}{1 + \exp(-\mathbf{u}_i^\top \bar{B} - \alpha)}\right]^{y_i} \left[\frac{1}{1 + \exp(\mathbf{u}_i^\top \bar{B} + \alpha)}\right]^{1-y_i},$$

where $\alpha$ is the intercept in logistic regression.

**Ordinal response:** We consider an ordinal logistic model for modeling the ordinal response. Suppose that there are $M$ ordered classes and $y_i \in \{1, \ldots, M\}$, then

$$\begin{aligned} P(y_i|\mathbf{u}_i, \Theta) &= \prod_{m=1}^{M} [\mathbf{P}(y_i \geq m|\mathbf{u}_i, \Theta) - \mathbf{P}(y_i \geq m + 1|\mathbf{u}_i, \Theta)]^{\mathbb{1}_{y_i = m}} \\ &= \prod_{m=1}^{M} \left[\frac{1}{1 + \exp(\mathbf{u}_i^\top \bar{B} + \alpha_m)} - \frac{1}{1 + \exp(\mathbf{u}_i^\top \bar{B} + \alpha_{m+1})}\right]^{\mathbb{1}_{y_i = m}}, \end{aligned}$$

where $\infty = \alpha_1 > \ldots > \alpha_M > \alpha_{M+1} = -\infty$ are the "cuts" along the direction $\mathbf{u}_i^\top \bar{B}$ that determines the probability of falling into each class, and $\mathbb{1}\{y_i = m\}$ is the indicator function for $y_i = m$.

**Multi-class categorical response:** We model the multi-class categorical response $y_i \in \{1, \ldots, M\}$ using the multinomial logistic regression. In this case $\bar{B} \in \mathbb{R}^{K \times M}$ is a matrix with $m^{th}$ column representing the coefficient for class $m$:

$$P(y_i|\mathbf{u}_i, \Theta) = \prod_{m=1}^{M} \left[ \frac{\exp(\mathbf{u}_i^\top \bar{B}_m + \alpha_m)}{\sum_{m'=1}^{M} \exp(\mathbf{u}_i^\top \bar{B}_{m'} + \alpha_{m'})} \right]^{\mathbb{1}\{y_i = m\}}$$

In Appendix B, we first derive the iterative parameter estimation procedures for the Gaussian data in B, and we then move to the non-Gaussian case in Appendix C.

#### A.2. Rank selection and weight tuning

In A.1, we described the SPEAR model with fixed weight parameter $w$ and rank $K$. Here, we give more details on our default choice of $w$ and $K$.

**Tuning the weight parameter $w$:** We tune the weight parameter adaptively from the data based on cross-validation on the response's deviance loss. To reduce the computational burden, we adopt the warm-start strategy: when running the model with a lower weight, we use model parameters from the higher weight before it as the starting point. Suppose that we divide the data into $T$ random folds $\cup_{t=1}^{T} \mathcal{D}_t$. Let $\hat{\Theta}(w; t)$ be the estimated posterior means for model parameters using data excluding fold $t$. Let $D(y_i, x_i|\hat{\Theta}(w; t))$ be the deviance loss evaluation at sample $i$ using the parameter $\hat{\Theta}(w; t)$, e.g, for Gaussian response, we have

$$D(y_i, x_i|\hat{\Theta}(w, t)) = \|y_i - [x_i^\top \hat{\boldsymbol{\beta}}(w, t)]\hat{\bar{\mathbf{B}}}(w, t)\|_2^2$$

where $\hat{\boldsymbol{\beta}}(w, t)$ and $\hat{\bar{\mathbf{B}}}(w, t)$ are the estimated posterior means for $\boldsymbol{\beta}$ and $\mathbf{B}$ using data excluding fold $t$ and at weight parameter $w$. Then, we can choose $w$ minimizing the average deviance loss:

$$D(Y, \mathbf{X}|\hat{\Theta}(w)) = \frac{1}{N} \sum_{t=1}^{T} \sum_{i \in \mathcal{D}_t} D(y_i, x_i|\hat{\Theta}(w, t)).$$

Let $w^*$ be the optimal weight minimizing the cross-validation error. In the 1 standard deviation rule, we choose the largest weight $w$ such that

$$D(Y, \mathbf{X}|\hat{\Theta}(w)) \leq D(Y, \mathbf{X}|\hat{\Theta}(w^*)) + \widehat{sd}\left(D(Y, \mathbf{X}|\hat{\Theta}(w^*))\right),$$

where $\widehat{sd}\left(D(Y, \mathbf{X}|\hat{\Theta}(w^*))\right)$ is the estiamted standard deviation of $D(Y, \mathbf{X}|\hat{\Theta}(w^*; t))$ by comparing the average deviance loss from different folds. We suggesting using $w^*$ when the goal is for better prediction and using the 1sd rule when more structure in $\mathbf{X}$ is preferred.

**Rank selection:** SPEAR chooses a weight parameter adaptively after fixing the rank $K$. When there is strong factor structure in $\mathbf{X}$ that is irrelevant to $Y$, choosing a rank $K$ too small can push our model towards small $w$, which can be unwanted in practice sometimes for interpretation. Here, we use recently developed statistical techniques and propose a simple novel approach as the default rank selection rule for SPEAR. We have assumed the following latent factor model for the features $\mathbf{X} \in \mathbb{R}^{N\times p}$:

$$\mathbf{X} = \mathbf{U}\mathbf{B} + \mathbf{E},$$

where $\mathbf{E} \in \mathbb{R}^{N\times p}$ is the unstructured noise matrix and $E_{ij} \overset{i.i.d}{\sim} N(0, \sigma_j^2)$. Without loss of generality, we always standardize $\mathbf{X}$ to have mean 0 and variance 1.

Now, let's regenerate a new noise matrix $\tilde{\mathbf{E}}$ where $\tilde{E}_{ij} \overset{i.i.d}{\sim} N(0, \sigma_j^2)$ follows the same distribution of $E_{ij}$. A key observation is that

$$\mathbf{X}^{\kappa} = \mathbf{U}\mathbf{B} + \mathbf{E} + \kappa\tilde{\mathbf{E}}$$

and

$$\mathbf{X}^{\frac{1}{\kappa}} = \mathbf{U}\mathbf{B} + \mathbf{E} - \frac{1}{\kappa}\tilde{\mathbf{E}}$$

are marginally independent if we consider both the randomness in $\mathbf{E}$ and the randomness in $\tilde{\mathbf{E}}$. This is a direct result of the Gaussianity and the fact that the covariance between $\mathbf{X}^{\kappa}$ and $\mathbf{X}^{\frac{1}{\kappa}}$:

$$cov(\mathbf{X}^{\kappa}, \mathbf{X}^{\frac{1}{\kappa}}) = \mathbf{0}.$$

Following this observation, we propose the following rank selection approach: This idea

**for** $\ell = 1, \ldots, L$ **do**

Generate $i.i.d$ Gaussian noise $\tilde{\mathbf{E}}$, and form $\mathbf{X}^{\kappa}$, $\mathbf{X}^{\frac{1}{\kappa}}$ with $\kappa = 0.1$.

Perform PCA on $\mathbf{X}^{\kappa}$, and let $\hat{\mathbf{X}}[k]$ be the reconstructed feature matrix with rank $k$ for $k = 1, \ldots, K_{\max}$.

Evaluate the reconstruction loss comparing $\hat{\mathbf{X}}[k]$ and $\mathbf{X}^{\frac{1}{\kappa}}$:

$$r_{kl} = \frac{1}{Np}\|\mathbf{X}^{\frac{1}{\kappa}} - \hat{\mathbf{X}}[k]\|_F^2$$

Select the rank with smallest average reconstruction loss

$$r_k = \frac{1}{L}\sum_{l=1}^{L} r_{kl}, \quad k^* = \arg\min_{k\in\{1,\ldots,K_{\max}\}} r_k$$

**end**

**Algorithm 1:** Rank selection in SPEAR

has been applied in the supervised regression problem for model selection (Guan and Tibshirani, 2020; Tian, 2020), but never to the rank selection.

In practice, we do not know the true noise level $\sigma_j^2$. Since we are most interested finding the strong factors, we can afford using a larger variance estimate. When replacing $\sigma_j^2$ by $\hat{\sigma}_j^2$ with $\hat{\sigma}_j^2 > \sigma_j^2$, this will introduce negative correlation between $\mathbf{X}^{\kappa}$ and $\mathbf{X}^{\frac{1}{\kappa}}$, which in turn favors smaller $K$. However, when the factor is strong, we expect them to be found even in this conservative setting. As a result, in the default SPEAR algorithm, instead of trying to estimate $\sigma_j^2$ accurately, we use an estimated upper bound $\hat{\sigma}_j^2 \in [0, 1]$, e.g., $\hat{\sigma}_j^2 = 1.0$ or 0.9. In our simulation and empirical studies, we have selected the model rank using this default strategy. We observe it works well for both cases: it selects the true ranks with high probability in simulations and ranks that do not push $w$ to 0 in real data experiments.

#### A.3. Automated selection of weight parameter ($w$) via cross-validation (CV)

SPEAR controls the level of prioritization in construction of predictive factors by adjusting the importance of the fit towards X through the weight parameter w. A low w (w≈0) will yield factors that emphasize prediction of Y, while higher values for w (w≥1) will construct factors that also can explain the structure of X.

We propose to automatically balance between the construction of factors and the prediction of the response through the weight parameter w, which is selected via cross-validation. SPEAR factors are constructed K times, where K is the number of folds used to split the training data, and the mean cross-validated error is calculated per weight w across folds. By default, the largest w whose mean cross-validated error falls within one standard deviation of the overall minimum cross-validated error is selected (Hastie et al., 2009), favoring higher values of w by design. Thus, SPEAR constructs predictive factors influenced by the multi-omics assays if data supported, returning a simple linear predictive model otherwise.

### B. Parameter estimation for Gaussian response

#### B.1. Review of the mean-field approximation

Let $\Theta = \{\theta_1, \dots, \theta_T\}$ be the collection of $T$-block of parameters, and $\mathcal{D}$ represent the data. Our goal is to approximate the posterior distribution of $\Theta$ with a simpler function $Q$ and find $Q$ minimizing their KL divergence between these two distributions:

$$KL(Q\|P) = \int Q(\Theta) \log \frac{Q(\Theta)}{P(\Theta|\mathcal{D})} d\Theta$$

Define the evidence lower bound (ELBO) to be the integral of the log ratio between the marginal density $P(\Theta, \mathcal{D})$ and $Q(\Theta)$:

$$ELBO(Q) = \int Q(\Theta) \log \frac{P(\Theta, P)}{Q(\Theta)} d\Theta = -KL(Q\|P) + \log P(\mathcal{D}) \tag{4}$$

The log marginal likelihood on the data $\log P(\mathcal{D})$ is called the evidence. It measures how well our model describes the data and is constant. Consequently, minimizing the KL-divergence between the variational distribution is equivalent of maximizing the ELBO. By choosing a proper variational distribution $Q$ such that integral of the log joint density or $\log Q$ over $Q$ is easy to calculate, we can find an optimal or local optimal solution to this maximization problem efficiently:

$$ELBO(Q) = \int Q(\Theta) \log P(\Theta, \mathcal{D}) - \int Q(\Theta) \log Q(\Theta)$$

$\mathcal{Q}$, the space of $Q$, often does not contain the actual posterior distribution $P(\Theta|\mathcal{D})$. However, variational Bayes has proven successful in many applied problems and researchers sometimes are willing to sacrifice accuracy for efficiency, especially in large-scale datasets.

One popular variational Bayesian method is mean field approximation, which fully decouples the parameters into several independent blocks and considers:

$$Q(\Theta) = \prod_{j=1}^{p} q_{\alpha_j}(\theta_j)$$

where $\alpha_j$ is the parameter determining the distribution $q_{\alpha_j}(\theta_j)$. We can often iteratively update $q_{\alpha_j}(\theta_j)$ in a computational efficient manner given the distributions of other parameters:

$$\begin{aligned}
&\int Q(\Theta)\left(\log P(\Theta,\mathcal{D}) - \log Q(\Theta)\right) \\
\propto &\int_j q_{\alpha_j}(\theta_j)\left(\left(\int_{\Theta_{-j}} Q_{-j}(\Theta_{-j})\log P(\Theta,\mathcal{D})d\Theta_{-j}\right) - \log q_{\alpha_j}(\theta_j)\right)d\theta_j
\end{aligned}$$

Fixing others, we choose the parameters $\alpha_j$ that improve most the ELBO:

$$L(Q) = \int Q(\Theta)\log\frac{P(Y,\Theta)}{Q(\Theta)} = E_q(\log P(Y|\Theta)) + \sum_{j=1}^{p}\left(E_{q_j}\log p(\theta_j) - E_{q_j}\log q(\theta_j)\right) \quad (5)$$

We can update the variational parameters $\alpha_1, \ldots, \alpha_p$ iteratively until the ELBO updates are deemed insignificant.

#### B.2. Variational distribution for SPEAR

Let $Q(\Theta \cup \boldsymbol{\mathcal{E}})$ be the mean field approximation to the posterior distribution of the model parameters. We assume $Q$ to be the product of a series of independent components:

$$\begin{aligned}
Q(\Theta \cup \boldsymbol{\mathcal{E}}) = &\left(\sum_{k=1}^{K}\prod_{j=1}^{p} Q(\beta_{jk})\right) \times \left(\sum_{k=1}^{K}\prod_{j=1}^{p} Q(B_{jk})\right) \times \left(\prod_{j=1}^{p_0} Q(\bar{B}_j)\right) \times \left(\prod_{i=1}^{n}\prod_{k=1}^{K} Q(\boldsymbol{\mathcal{E}}_{ik})\right) \\
&\times \left(\prod_{j=1}^{p} Q(\nu_j)\right) \times \left(\prod_{j=1}^{p_0} Q(\bar{\nu}_j)\right) \times \left(\prod_{g=1}^{G}\prod_{k=1}^{K} Q(\tau_{gk})\right) \times \left(\prod_{g=1}^{G}\prod_{k=1}^{K} Q(\pi_{gk})\right)
\end{aligned}$$

The specific forms for each component are given below.

$$\begin{aligned}
&Q(\bar{B}_{jk}) = \mathcal{N}(\bar{B}_{jk}|\bar{\mu}_{jk}, \bar{\zeta}^2_{jk}),\ Q(\boldsymbol{\mathcal{E}}_{ik}) = \mathcal{N}(\boldsymbol{\mathcal{E}}_{ik}|0, e_{ik}), \\
&Q(\nu_j) = \Gamma(\nu_j|\tilde{a}^X_{j0}, \tilde{c}^X_{j0}),\ Q(\bar{\nu}_j) = \Gamma(\nu_j|\tilde{a}^Y_{j0}, \tilde{c}^Y_{j0}), \\
&Q(\tau_{gk}) = \Gamma(\tau_{gk}|\tilde{a}_{gk1}, \tilde{c}_{gk1}),\ Q(\pi_{gk}) = \Gamma(\pi_{gk}|\tilde{a}_{gk2}, \tilde{c}_{gk2}),
\end{aligned}$$

and also,

$$Q(\beta_{jk}) \coloneqq Q(\hat{\beta}_{jk}, \gamma_{jk}) = \begin{cases} \mathcal{N}(\hat{\beta}_{jk}|\mu_{jk}, \zeta^2_{jk}) & \text{if} \quad \gamma_{jk} = 1,\ (w.p\ \omega_{jk}) \\ \mathcal{N}(\hat{\beta}_{jk}|0, \zeta^2_{0jk}) & \text{otherwise},\ (w.p\ 1-\omega_{jk}) \end{cases},$$

$$Q(B_{jk}) \coloneqq Q(\hat{B}_{jk}, s_{jk}) = \begin{cases} \mathcal{N}(\hat{B}_{jk}|\tilde{\mu}_{jk}, \tilde{\zeta}^2_{jk}) & \text{if} \quad s_{jk} = 1\ (w.p\ \tilde{\omega}_{jk}) \\ \mathcal{N}(\hat{B}_{jk}|0, \tilde{\zeta}^2_{0jk}) & \text{otherwise},\ (w.p\ 1-\tilde{\omega}_{jk}) \end{cases}.$$

#### B.3. Formula for different expectations

We use $\langle . \rangle$ to represent the expectation under the variational distribution. Then,

- $\langle \beta_{jk} \rangle = \langle \gamma_{jk} \rangle \langle \hat{\beta}_{jk} \rangle = \omega_{jk}\mu_{jk}$, $\langle \beta_{jk}^2 \rangle = \omega_{jk}\left(\mu_{jk}^2 + \zeta_{jk}^2\right)$, $\langle \hat{\beta}_{jk}^2 \rangle = \langle \beta_{jk}^2 \rangle + (1-\omega_{jk})\zeta_{0jk}^2$.
- $\langle B_{jk} \rangle = \langle s_{jk} \rangle \langle \hat{B}_{jk} \rangle = \tilde{\omega}_{jk}\tilde{\mu}_{jk}$, $\langle B_{jk}^2 \rangle = \tilde{\omega}_{jk}\left(\tilde{\mu}_{jk}^2 + \tilde{\zeta}_{jk}^2\right)$, $\langle \hat{B}_{jk}^2 \rangle = \langle B_{jk}^2 \rangle + (1-\tilde{\omega}_{jk})\tilde{\zeta}_{0jk}^2$.
- $\langle \boldsymbol{\mathcal{E}}_{ik} \rangle = 0$, $\langle \boldsymbol{\mathcal{E}}_{ik}^2 \rangle = e_{ik}$.
- $\langle u_{ik} \rangle = z_i^{\perp} \langle \boldsymbol{\beta}_k \rangle$, $\langle u_{ik}^2 \rangle = (z_i^{\perp} \langle \boldsymbol{\beta}_k \rangle)^2 + \sum_{j=1}^p z_{ij}^2 (\langle \boldsymbol{\beta}_k^2 \rangle - \langle \boldsymbol{\beta}_k \rangle^2) + \langle \boldsymbol{\mathcal{E}}_{ik}^2 \rangle$.

We collect some results about the Gamma and Beta distributions below. Let $x$ be a variable with $\Gamma(x|a,c)$ or $\text{Beta}(x|a,c)$ depending on the context. Let $\Psi(a,c) = \frac{d \log \Gamma(a,c)}{da}$ be the first-order polyGamma distribution, then,

$$\mathbb{E}_{\Gamma}[x] = \frac{a}{c},\ \mathbb{E}_{\Gamma}[\log x] = \log c + \Psi(a,c),$$
$$\mathbb{E}_{\text{Beta}}[x] = \frac{a}{a+c},\ \mathbb{E}_{\text{Beta}}[\log x] = \Psi(a+c,1) - \Psi(a,1),\ \mathbb{E}_{\text{Beta}}[\log(1-x)] = \Psi(a+c,1) - \Psi(c,1).$$

We can get the expectations $\langle \nu_j \rangle$, $\langle \bar{\nu}_j \rangle$, $\langle \tau_{gk} \rangle$, $\langle \log \tau_{gk} \rangle$, $\langle \pi_{gk} \rangle$, $\langle \log \pi_{gk} \rangle$, $\langle \log(1 - \pi_{gk}) \rangle$ straightforwardly based on the above formulas.

#### B.4. Parameter update

**Update** $Q(\beta_{jk})$**.** We let $L_0(\beta_{jk})$ denote the expected log prior on $\beta$, $L_1(\beta_{jk})$ denote the expected log data density (relevant to $\beta$) and $L_2(\beta_{jk})$ be expected log variational distribution. (We use such notations throughout this supplement, e.g., for any parameter $\theta$, we use $L_0(\theta), L_1(\theta), L_2(\theta)$ to denote the logarithms on the part relevant to prior, data density and variational distribution.) Then,

$$L_0(\beta_{jk}) \propto -\frac{1}{2}\langle \hat{\beta}_{jk}^2 \rangle \langle \tau_{g_j k} \rangle + \langle \gamma_{jk} \rangle \langle \ln \pi_{g_j \pi} \rangle + \langle \gamma_{jk} \rangle \langle \ln(1 - \pi_{g_j \pi}) \rangle, \tag{6}$$

$$L_1(\beta_{jk}) \propto -\frac{1}{2}\sum_{j'=1}^{p} w_{j'} \langle \nu_{j'} \rangle \langle \|\mathbf{X}_{j'} - \mathbf{U}\mathbf{B}_{j'}\|_2^2 \rangle - \frac{1}{2}\sum_{j'=1}^{p_0} \langle \bar{\nu}_{j'} \rangle \langle \|\mathbf{Y}_{j'} - \mathbf{U}\bar{\mathbf{B}}_{j'}\|_2^2 \rangle, \tag{7}$$

$$L_2(\beta_{jk}) \propto -\omega_{jk}\frac{1 + \ln \zeta_{jk}^2}{2} - (1-\omega_{jk})\frac{1 + \ln \zeta_{0jk}^2}{2} + (1-\omega_{jk})\ln(1-\omega_{jk}) + \omega_{jk}\ln \omega_{jk}. \tag{8}$$

Re-arrange the terms for $L_1(\beta_{jk})$, we have

$$L_1(\beta_{jk}) \propto \sum_{i=1}^{n} (I_{i1} - I_{i2})\, z_{ij} \langle \beta_{jk} \rangle - \frac{1}{2}\sum_{i=1}^{n} I_{i3} z_{ij}^2 \langle \beta_{jk}^2 \rangle \tag{9}$$

where

$$I_{i1} = \sum_{j'=1}^{p} \langle \nu_{j'} \rangle w_{j'} \left\langle x_{ij'} - \sum_{k' \neq k} u_{ik'} B_{j'k'} \right\rangle \langle B_{j'k} \rangle + \sum_{j'=1}^{p_0} \langle \bar{\nu}_{j'} \rangle \left\langle y_{ij'} - \sum_{k' \neq k} u_{ik'} \bar{B}_{j'k'} \right\rangle \langle \bar{B}_{j'k} \rangle$$

$$I_{i2} = \sum_{j'=1}^{p} \langle \nu_{j'} \rangle w_{j'} \left\langle u_{ik}^{\setminus j} \right\rangle \left\langle B_{j'k}^2 \right\rangle + \sum_{j'=1}^{p_0} \langle \bar{\nu}_{j'} \rangle \left\langle u_{ik}^{\setminus j} \right\rangle \left\langle \bar{B}_{j'k}^2 \right\rangle,$$

$$I_{i3} = \sum_{j'=1}^{p} \langle \nu_{j'} \rangle w_{j'} \left\langle B_{j'k}^2 \right\rangle + \sum_{j'=1}^{p_0} \langle \bar{\nu}_{j'} \rangle \left\langle \bar{B}_{j'k}^2 \right\rangle,$$

where $u_{ik}^{/j} = \sum_{j' \neq j} z_{ij'} \beta_{j'k}$ is factor $k$ constructed excluding the $j^{th}$ feature. It is easy to check that

$$\langle \gamma_{jk} \rangle = \omega_{jk},\ \langle \hat{\beta}_{jk} \rangle = \mu_{jk}\omega_{jk},\ \langle \hat{\beta}_{jk}^2 \rangle = \left(\mu_{jk}^2 + \zeta_{jk}^2\right)\omega_{jk} + (1-\omega_{jk})\zeta_{0jk}^2,$$
$$\langle \beta_{jk} \rangle = \mu_{jk}\omega_{jk},\ \langle \beta_{jk}^2 \rangle = \left(\mu_{jk}^2 + \zeta_{jk}^2\right)\omega_{jk}.$$

The variational parameters $(\mu_{jk}, \zeta_{jk}^2, \zeta_{0jk}^2)$ are chosen to maximize the ELBO, or equivalently, maximize

$$L(\beta_{jk}) = L_0(\beta_{jk}) + L_1(\beta_{jk}) - L_2(\beta_{jk}).$$

Omitting the part irrelevant to $\beta_{jk}$, and letting $A_1 = \sum_i (I_{i1} - I_{i2}) z_{ij}$, $A_2 = \sum_i I_{i3} z_{ij}^2$, we have

$$\begin{aligned} L(\beta_{jk}) = {} & \omega_{jk} \left\{ A_1 \mu_{jk} - \frac{1}{2}(A_2 + \langle \tau_{g_j k} \rangle)\left(\mu_{jk}^2 + \zeta_{jk}^2\right) + \frac{1}{2} \ln \zeta_{jk}^2 \right\} \\ & + (1-\omega_{jk}) \left\{ -\frac{1}{2} \langle \tau_{g_j k} \rangle \zeta_{0jk}^2 + \frac{1}{2} \ln \zeta_{0jk}^2 \right\} \\ & + \omega_{jk} \langle \ln \frac{\pi_{g_j k}}{1 - \pi_{g_j k}} \rangle - \omega_{jk} \ln \frac{\omega_{jk}}{1 - \omega_{jk}} - \ln(1 - \omega_{jk}) \end{aligned} \tag{10}$$

- $\mu_{jk}$ is chosen as

$$\mu_{jk} = \arg\max \left( A_1 \mu_{ij} - \frac{1}{2} A_2 \mu_{ij}^2 - \frac{\langle \tau_{g_j k} \rangle}{2} \mu_{jk}^2 \right) = \frac{A_1}{\langle \tau_{g_j k} \rangle + A_2}.$$

- $\zeta_{jk}^2$ and $\zeta_{0jk}^2$ are chosen as

$$\zeta_{jk}^2 = \arg\max \left( -\frac{1}{2} A_2 \zeta_{jk}^2 - \frac{\langle \tau_{g_j k} \rangle}{2} + \frac{1}{2} \ln \zeta_{jk}^2 \right) = \frac{1}{\langle \tau_{g_j k} \rangle + A_2},$$
$$\zeta_{0jk}^2 = \arg\max \left( -\frac{\langle \tau_{g_j k} \rangle}{2} + \frac{1}{2} \ln \zeta_{jk}^2 \right) = \frac{1}{\langle \tau_{g_j k} \rangle}.$$

- $\omega_{jk}$ is chosen as

$$\begin{aligned} \omega_{jk} &= \arg\max \left( \omega_{jk} \left( \frac{\mu_{jk}^2}{2\zeta_{jk}^2} + \frac{1}{2} \ln \frac{\zeta_{jk}^2}{\zeta_{0jk}^2} + \ln \langle \frac{\pi_{g_j k}}{1 - \pi_{g_j k}} \rangle \right) - \omega_{jk} \ln \omega_{jk} - (1 - \omega_{jk}) \log(1 - \omega_{jk}) \right) \\ &= \frac{\exp(\lambda_{jk})}{1 + \exp(\lambda_{jk})} \end{aligned}$$

where $\lambda_{jk} = \frac{\mu_{jk}^2}{2\zeta_{jk}^2} + \frac{1}{2} \ln \frac{\zeta_{jk}^2}{\zeta_{0jk}^2} + \ln \langle \frac{\pi_{g_j k}}{1 - \pi_{g_j k}} \rangle$.

**Update $Q(B_{jk})$.** Updating $B_{jk}$, $L_0(B_{jk})$ and $L_2(B_{jk})$ is similar to that of $L_0(\beta_{jk})$ and $L_2(\beta_{jk})$, except for replacing all quantities by their corresponding component for $B_{jk}$. The data density related part is

$$L_1(B_{jk}) \propto -\frac{1}{2} w_j \langle \nu_j \rangle \langle \|\mathbf{X}_j - \mathbf{U}\mathbf{B}_j\|_2^2 \rangle. \tag{11}$$

Rearrange the terms, we have

$$L_1(B_{jk}) \propto w_j \langle \nu_j \rangle \left( \langle \mathbf{X}_j - \mathbf{U}_{k^c} \mathbf{B}_{k^c,j} \rangle^{\perp} \langle \mathbf{U}_k \rangle \langle B_{jk} \rangle - \frac{1}{2} \langle \|\mathbf{U}_k\|_2^2 \rangle \langle B_{jk}^2 \rangle \right).$$

We set $A_1 = w_j \langle \nu_j \rangle \langle \mathbf{X}_j - \mathbf{U}_{k^c} \mathbf{B}_{k^c,j} \rangle^{\perp} \langle \mathbf{U}_k \rangle$ and $A_2 = w_j \langle \nu_j \rangle \langle \|\mathbf{U}_k\|_2^2 \rangle$, then,

$$\begin{aligned} L(B_{jk}) = {} & \tilde{\omega}_{jk} \left\{ A_1 \tilde{\mu}_{jk} - \frac{1}{2}(A_2 + \langle \tau_{g_j k} \rangle) \left( \tilde{\mu}_{jk}^2 + \tilde{\zeta}_{jk}^2 \right) + \frac{1}{2} \ln \tilde{\zeta}_{jk}^2 \right\} \\ & + (1 - \tilde{\omega}_{jk}) \left\{ -\frac{1}{2} \langle \tau_{g_j k} \rangle \tilde{\zeta}_{0jk}^2 + \frac{1}{2} \ln \tilde{\zeta}_{0jk}^2 \right\} \\ & + \tilde{\omega}_{jk} \langle \ln \frac{\pi_{g_j k}}{1 - \pi_{g_j k}} \rangle - \tilde{\omega}_{jk} \ln \frac{\tilde{\omega}_{jk}}{1 - \tilde{\omega}_{jk}} - \ln(1 - \tilde{\omega}_{jk}) \end{aligned} \tag{12}$$

Hence, our update is

$$\begin{aligned} \tilde{\zeta}_{jk}^2 &= \frac{1}{A_2 + \langle \tau_{g_j k} \rangle}, \ \tilde{\zeta}_{0jk} = \frac{1}{\langle \tau_{g_j k} \rangle}, \\ \tilde{\mu}_{jk} &= \frac{A_1}{A_2 + \langle \tau_{g_j k} \rangle}, \ \tilde{\omega}_{jk} = \frac{\exp(\lambda_{jk})}{1 + \exp(\lambda_{jk})}, \end{aligned}$$

where $\lambda_{jk} = \frac{\tilde{\mu}_{jk}^2}{2\tilde{\zeta}_{jk}^2} + \frac{1}{2} \ln \frac{\tilde{\zeta}_{jk}^2}{\tilde{\zeta}_{0jk}^2} + \ln \langle \frac{\pi_{g_j k}}{1 - \pi_{g_j k}} \rangle$.

**Update $Q(\bar{B}_j)$.** We set $A_1 = \langle \bar{\nu}_j \rangle \langle \mathbf{U} \rangle^{\perp} \langle \mathbf{Y}_j \rangle$ and $A_2 = \langle \bar{\nu}_j \rangle \langle \mathbf{U}^{\perp} \mathbf{U} \rangle + \Lambda$, where $\Lambda = diag\{\tau_0^2, \ldots, \tau_0^2\}$, the ELBO related to $Q(\bar{B}_{jk})$ is

$$L(\overline{\mathbf{B}}_j) \propto A_1^{\perp} \langle \overline{\mathbf{B}}_j \rangle - \frac{1}{2} \langle \overline{\mathbf{B}}_j^{\perp} A_2 \overline{\mathbf{B}}_j \rangle + \frac{1}{2} \sum_{k=1}^{K} \ln \bar{\zeta}_{jk}^2 = A_1^{\perp} \bar{\mu}_j - \frac{1}{2} \bar{\mu}_j^{\top} A_2 \bar{\mu}_j - \frac{1}{2} \sum_{k=1}^{K} A_{2kk} \xi_{jk}^2 + \frac{1}{2} \sum_{k=1}^{K} \ln \bar{\zeta}_{jk}^2.$$

The optimal updating parameters is

$$\bar{\mu}_j = A_2^{-1} A_1, \ \bar{\zeta}_{jk}^2 = \frac{1}{A_{2kk}}.$$

We will now discuss the constrained solution where we require $\|\bar{\mu}_j\|_2 \geq L_{\overline{B}_j}$. This extra constraint is to alleviate the extra volatility due to the oscillation between $\overline{B}$ and $\boldsymbol{\beta}$ when $w$, the weight on $\mathbf{X}$, is small. By default, we let $L_{\overline{B}_j} = (1 - w) \vee 0$. This constraint does not affect $\bar{\zeta}_{jk}^2$, but we no longer has a closed form expression for $\bar{\mu}_j$. However, we can numerically find the optimal solution:

1. Consider the ridge penalized problem for $\bar{\mu}_j$:

$$\bar{\mu}_j(\alpha) = \arg\max_{\bar{\mu}_j} A_1^{\top} \bar{\mu}_j - \frac{1}{2} \bar{\mu}_j^{\top} (A_2 + \alpha \mathbf{Id}) \bar{\mu}_j$$

2. Let $\alpha^*$ be the largest non-positive value such that $\|\bar{\mu}_j(\alpha^*)\|_2^2 \geq L_{\overline{B}_j}$, which can be found via binary search.

3. Then, $\bar{\mu}_j = \bar{\mu}_j(\alpha^*)$ solves the constrained problem (Guan, 2021; Guan, 2022).

$$\bar{\mu}_j(\alpha) = \arg \max_{\bar{\mu}_j : \|\bar{\mu}_j\|_2^2 \geq L_{\overline{B}_j}} A_1^\top \bar{\mu}_j - \frac{1}{2} \bar{\mu}_j^\top A_2 \bar{\mu}_j.$$

**Update $Q(\boldsymbol{\mathcal{E}})$.** The expected logarithm from the ELBO calculation related to $\boldsymbol{\mathcal{E}}_{ik}$ is given below (used the fact that $\langle \boldsymbol{\mathcal{E}}_{ik} \rangle = 0$):

$$L(\boldsymbol{\mathcal{E}}_{ik}) \propto \frac{1}{2} \left( -\sum_{j=1}^{p} w_j \langle \nu_j \rangle \langle B_{ij}^2 \rangle e_{jk}^2 - \sum_{j=1}^{p_0} \langle \bar{\nu}_j \rangle \langle \bar{B}_{ij}^2 \rangle e_{jk}^2 - \frac{e_{ik}^2}{2} + \ln e_{ik}^2 \right).$$

To avoid instability, we also require $e_{jk}^2 \geq L_{\boldsymbol{\mathcal{E}}}$ and $L_{\boldsymbol{\mathcal{E}}} = 0.1$ by default. Let $A = \frac{1}{\sum_{j=1}^{p} w_j \langle \nu_j \rangle \langle B_{jk}^2 \rangle + \sum_{j=1}^{p_0} \langle \bar{\nu}_j \rangle \langle \bar{B}_{jk}^2 \rangle e_{ik}^2 + 1}$. The optimal solution under this constraint is

$$e_{jk}^2 = \begin{cases} A & \text{if } A \geq L_{\boldsymbol{\mathcal{E}}} \\ L_{\boldsymbol{\mathcal{E}}} & \text{otherwise} \end{cases}.$$

**Update $Q(\tau)$.** The parameter $\tau$ appear only at the prior and the hyper prior, its related ELBO at parameter $\theta = (\tilde{a}_{gk}^1, \tilde{c}_{gk}^1)$ is

$$L(\tau_{gk}) \propto -\left( \frac{\sum_{g_j = g} \left( \langle \hat{\beta}_{jk}^2 \rangle + \langle \hat{B}_{jk}^2 \rangle \right)}{2} + c_1 - \tilde{c}_{gk}^1 \right) \langle \tau_{gk} \rangle + \left( |\mathcal{G}_g| + a_1 - \tilde{a}_{gk} \right) \langle \ln \tau_{gk} \rangle + \ln \Gamma(\tilde{a}_{gk}^1, \tilde{c}_{gk}^1),$$

where $\mathcal{G}_g$ is the set of feature indices in group $g$ and $|\mathcal{G}_g|$ is the set size. The optimal update rule is

$$\tilde{a}_{\ell k}^1 = a_0 + |\mathcal{G}_\ell|, \quad \tilde{c}_{\ell k}^1 = c_1 + \frac{\sum_{g_j = \ell} \left( \langle \hat{\beta}_{jk}^2 \rangle + \langle \hat{B}_{jk}^2 \rangle \right)}{2}.$$

Intuitively, when $\langle \tau_{\ell k} \rangle = \frac{\tilde{a}_{\ell k}^1}{\tilde{b}_{\ell k}}$ is large, we will have a prior more concentrated around 0, and hence, a denser model with small effects. When $\langle \tau_{\ell k} \rangle$ is very small, on the other hand, we will only let a feature being non-zero if it's effect size is very large. As a result, the model does not favor $\langle \tau_{\ell k} \rangle$ being too small or too large. Since $\tilde{a}_{\ell k}^1 = a_0 + |\mathcal{G}_\ell|$ is fixed, if we want $\langle \tau_{\ell k} \rangle \in [L_{low}, L_{up}]$, we can find an optimal $\tilde{c}_\ell^1$ in the range $[\frac{\tilde{a}_{\ell k}^1}{L_{up}}, \frac{\tilde{a}_{\ell k}^1}{L_{low}}]$ (that is left/right censored at $\frac{\tilde{a}_{\ell k}^1}{L_{up}}$ and $\frac{\tilde{a}_{\ell k}^1}{L_{low}}$). By default, we let $L_{low} = \frac{\ln p}{n}$ and $L_{up} = 1$.

**Update $Q(\pi)$.** The ELBO related to $\pi_{\ell k}$ at the variational parameter $\theta = (\tilde{a}_{\ell k}^2, \tilde{c}_{\ell k}^2)$ is

$$L(\pi_{\ell k}) \propto \left( a_2 + \sum_{g_j = \ell} \left( \langle \gamma_{jk} \rangle + \langle s_{jk} \rangle \right) - \tilde{a}_{\ell k}^2 \right) \langle \ln \pi_{\ell k} \rangle + \left( c + \sum_{g_j = \ell} \left( 2 - \langle \gamma_{jk} \rangle - \langle s_{jk} \rangle \right) - \tilde{c}_{\ell k}^2 \right) \langle \ln(1 - \pi_{\ell k}) \rangle + \ln \text{Beta}(\tilde{a}_{\ell k}^2, \tilde{c}_{\ell k}^2).$$

The optimal solution is

$$\tilde{a}_{\ell k}^2 = a_2 + \sum_{g_j=\ell} \left(\langle\gamma_{jk}\rangle + \langle s_{jk}\rangle\right), \ \tilde{c}_{\ell k}^2 = c + \sum_{g_j=\ell} \left(2 - \langle\gamma_{jk}\rangle - \langle s_{jk}\rangle\right).$$

The average sparsity level is $\langle\pi_{\ell k}\rangle = \frac{\tilde{a}_{\ell k}^2}{\tilde{a}_{\ell k}^2+\tilde{c}_{\ell k}^2} = \frac{\tilde{a}_{\ell k}^2}{a_2+c_2+2|\mathcal{G}_\ell|}$. We can also have a user specified upper bound on this average sparsity level. Let $\alpha$ be the specified sparsity level upper bound with a default value at 0.5. We find the optimal solution under the constraint

$$\tilde{c}_{\ell k}^2 + \tilde{a}_{\ell k}^2 = a_2 + c_2 + 2|\mathcal{G}_\ell|, \tilde{a}_{\ell k}^2 \le \alpha \left(a_2 + c_2 + 2|\mathcal{G}_\ell|\right).$$

If the optimal solution already satisfies the constraint, we don't need any modifications; otherwise, we let

$$\tilde{a}_{\ell k}^2 = \alpha \left(a_2 + c_2 + 2|\mathcal{G}_\ell|\right), \ \tilde{a}_{\ell k}^2 = (1-\alpha)\left(a_2 + c_2 + 2|\mathcal{G}_\ell|\right).$$

**Update $Q(\nu)$ and $Q(\bar{\nu})$.** Let $A_1 = w_j \left(\frac{N_j}{2} + a_0\right)$, $A_2 = w_j \left(\frac{\langle\|\mathbf{X}_j - \mathbf{U}\mathbf{B}_k\|_2^2\rangle}{2} + c_0\right)$, the ELBO related to $\nu_j$ is

$$L(\nu_j) \propto \langle \ln \Gamma(\nu_j|A_1, A_2)\rangle - \langle \ln \Gamma(\nu_j|\tilde{a}_j^X, \tilde{c}_j^Y)\rangle.$$

The optimal solution is

$$\tilde{a}_j^X = w_j \left(\frac{N_j}{2} + a_0\right), \ \tilde{c}_j^Y = w_j \left(\frac{\langle\|\mathbf{X}_j - \mathbf{U}\mathbf{B}_j\|_2^2\rangle}{2} + c_2\right).$$

Let $A_1 = \left(\frac{N_j}{2} + a_0\right)$, $A_2 = \left(\frac{\langle\|\mathbf{Y}_j - \mathbf{U}\mathbf{B}_k\|_2^2\rangle}{2} + c_0\right)$, the ELBO related to $\bar{\nu}_j$ is

$$L(\bar{\nu}_j) \propto \langle \ln \Gamma(\bar{\nu}_j|A_1, A_2)\rangle - \langle \ln \Gamma(\bar{\nu}_j|\tilde{a}_j^Y, \tilde{c}_j^Y)\rangle.$$

The optimal solution is

$$\tilde{a}_j^Y = \left(\frac{N_j}{2} + a_0\right), \ \tilde{c}_j^Y = \left(\frac{\langle\|\mathbf{Y}_j - \mathbf{U}\mathbf{B}_j\|_2^2\rangle}{2} + c_0\right).$$

**ELBO increase.** We can keep track of the ELBO increase by adding up the ELBO increase in each updating step. For given parameters, e.g., $\beta_{jk}$, let $\theta^{old}$ be the values for the variational parameters that are currently under investigation, and let $\theta$ be the updated values. We can calculate the related ELBO increase as follows:

$$\Delta(\beta_{jk}) = L(\beta_{jk}) - L_{old}(\beta_{jk}),$$

where $L(\beta_{jk})$ is the ELBO related to $\beta_{jk}$ using $\theta$ and $L_{old}(\beta_{jk})$ is the ELBO related to $\beta_{jk}$ using $\theta^{old}$. The total increase in ELBO can be calculated as

$$\begin{aligned}\Delta = &\sum_{k=1}^{K}\sum_{j=1}^{p} \Delta(\beta_{jk}) + \sum_{k=1}^{K}\sum_{j=1}^{p} \Delta(B_{jk}) + \sum_{j=1}^{\bar{p}} \Delta(\bar{B}_j) + \sum_{j=1}^{p} \Delta(\nu_j) + \sum_{j=1}^{\bar{p}} \Delta(\bar{\nu}_j) \\ &+ \sum_{k=1}^{K}\sum_{\ell=1}^{M} \left(\Delta(\tau_{\ell k}) + \Delta(\pi_{\ell k})\right).\end{aligned}$$

#### C. Parameter estimation with non-Gaussian response

Currently, SPEAR also supports two-class or multi-class classifications and ordinal regression. For these response types, the evidence lower bound under the given variational distribution becomes intractable. There are usually two ways for updating when the expectation is intractable: (1) find a lower bound of the ELBO that has closed form and optimize for this lower bound, or (2) use a numerical estimate of the gradient and update using gradient descent. There are pros and cons for both methods. For the former, we no longer optimize with respect to the original objective. For the later, there is variability of the derivative as well as the objective value due to the stochastic nature, and this can incur a large computational cost in order to have low variability. Here, we consider the approach of using the lower bound. The lower bounds for all three data types are based on results from **?**. The bound from **?** is used for the logistic model, and an variant is used for bounding the multinomial logistic regression loss **?**. We further extend this result to ordinal regression in this paper.

##### C.1. Lower Bounds of ELBO

If the expectation of the log-likelihood under the variational distribution is intractable, one popular way to circumvent it is to consider a good lower bound of this expectation that has a nice form with the help from extra augmenting variables and optimize in the enlarged parameter space. In general, let $f(\theta)$ be the log likelihood function. If its expectation is intractable under the variational distributions, but the quadratic function of $\theta$ can be explicitly calculated. Then, if there exist functions $a(\xi)$ and $A(\xi)$ for the augmenting variable $\xi$, such that

$$f(\theta) \geq a(\xi)^T\theta + \theta^T A(\xi)\theta + C(\xi) := g(\xi, \theta)$$

We have $f(\theta) \geq \max_\xi[f(\xi) + a(\xi)^T\theta + \theta^T A(\xi)\theta + C(\xi)]$, and

$$E_q f(\theta) \geq \max_\xi [C(\xi) + E_q\theta^T A(\xi)\theta + E_q(a(\xi)^T\theta)].$$

Hence, we can optimize for $q$ given $\xi$, and optimize for $\xi$ given $q$. This guarantees that the augmented lower-bound function $E_q g(\xi, \theta)$ is non-decreasing in the updating steps. We will consider two particular response types: logistic regression (two classes or multiple classes) and ordinal regression. The key results used to derive these lower bounds are from **?**: for any $\eta$, let $f(\eta) = \frac{1}{\exp(-\eta)+1}$ and $\lambda(\eta) = \tanh(\frac{\eta}{2})/(4\eta)$, then, we have

$$f(\eta) \geq f(\xi)\exp\left(\frac{\eta - \xi}{2} - \lambda(\xi)(\eta^2 - \xi^2)\right),\ \forall\xi. \tag{13}$$

A slightly more relaxed bound is

$$f(\eta) \geq f(\xi)\exp\left(\frac{\eta - \xi}{2} - \lambda(\xi)(\eta^2 - \xi^2) - \epsilon\eta^2\right),\ \forall\xi, \forall\epsilon \geq 0 \tag{14}$$

We let $\epsilon \geq 0$ be a user specified small constant, and it can encourage more numerical stability in the case of perfect separation when being positive.

##### C.2. Two-class logistic regression

Let $\eta_i = (\mathbf{u}_i^T\mathbf{B}_j + \alpha)$ be our link function value for response $j$ and sample $i$ in the two-class logistic model, and $\alpha$ is the intercept. Let $H_i = (2S_i - 1)\eta_i$ where $S_i \in \{0, 1\}$

indicates the class label. The expected log likelihood of sample $i$, $\langle \ell_i \rangle = \langle \log \frac{1}{1+\exp(\eta_i)} \rangle$, can be lower bounded with (14):

$$\langle \ell_i \rangle \geq \max_{\xi_i} \frac{\langle H_i \rangle - \xi_i}{2} - \lambda(\xi_i)(\langle H_i^2 \rangle - \xi_i^2) - \epsilon \langle H_i^2 \rangle.$$

Following the same argument as in **?**, despite the additional term $\epsilon$, the choice maximizing the lower bound given the variational distribution is

$$\xi_i = \sqrt{\langle H_i^2 \rangle} = \sqrt{\langle \eta_i^2 \rangle}. \tag{15}$$

Given $\xi$, we approximate the logistic with a Gaussian, and we update the variational distribution considering the lower bound

$$\underline{\ell}_i = -\left(\lambda(\xi_i) + \epsilon\right) \left\{ \left( \eta_i - \frac{(2S_i - 1)}{4(\lambda(\xi_i) + \epsilon)} \right)^2 \right\} \tag{16}$$

In other words, we are equivalently considering the following response when updating the variational distribution:

$$\underline{Y}_i \sim \mathcal{N}\left(\frac{(S_i - \frac{1}{2})}{2(\lambda(\xi_i) + \epsilon)} - \alpha, \frac{1}{2(\lambda(\xi_i) + \epsilon)}\right).$$

where the coefficient $\alpha$ can be first estimated by

$$\alpha = \frac{\sum_i \left( (\lambda(\xi_i) + \epsilon) \left( \underline{Y}_i - \langle \mathbf{u}_i^T \mathbf{B}_j \rangle \right) \right)}{2 \sum_i \left( \lambda(\xi_i) + \epsilon \right)}.$$

#### C.3. Ordinal logistic regression

Suppose that we are looking at the response $Y_i$ that takes value in $\{1, \ldots, M\}$. 1 is the base class and we are interested in estimation $P(Y_i \geq m)$ for $m = 2, \ldots, M$. Let $\eta_i = \mathbf{u}_i^T \mathbf{B}_j$ and $\infty = \alpha_1 > \ldots > \alpha_K > \alpha_{K+1} = -\infty$. The probability for observing a label at least $k$ is modeled as $P(Y_i \geq m) = \frac{1}{1+\exp(-\eta_i - \alpha_m)}$. The log-likelihood contributed by sample $i$ is

$$\ell_i = \sum_{m=1}^{M} \mathbb{1}(Y_i = m) \log \left( P(Y_i \geq m) - P(Y_i \geq m+1) \right).$$

After some simple rearrangements, we have

$$\ell_i = \begin{cases} -\eta_i - \alpha_2 - \log(1 + \exp(-\eta_i - \alpha_2)) & \text{if } Y_i = 1 \\ -\log(1 + \exp(-\eta_i - \alpha_M)) & \text{if } Y_i = M \\ -\eta_i + \log(\exp(-\alpha_{y_i+1}) - \exp(-\alpha_{y_i})) - \log(1 + \exp(-\eta_i - \alpha_{y_i})) - \log(1 + \exp(-\eta_i - \alpha_{y_i+1})) & \text{otherwise} \end{cases}.$$

We now apply (14) to each of the terms, and introduce auxiliary variables $\xi_{im}$ for $m = 2, \ldots, M$. The lower bound of the expected log likelihood given $\xi_{im}$ and $\alpha_m$ is then:

$$\langle \underline{\ell}_i \rangle = \begin{cases} \langle -\frac{\eta_i}{2} - \left(\lambda(\xi_{i,2}) + \epsilon\right) \left(\eta_i^2 + 2\alpha_2 \eta_i\right) \rangle & \text{if } y_i = 1 \\ \langle \frac{\eta_i}{2} - \left(\lambda(\xi_{iM}) + \epsilon\right) \left(\eta_i^2 + 2\alpha_M \eta_i\right) \rangle & \text{if } y_i = M \\ \langle -\left(\lambda(\xi_{i,y_i}) + \epsilon\right) \left(\eta_i^2 + 2\alpha_{y_i} \eta_i\right) - \left(\lambda(\xi_{i,y_i+1}) + \epsilon\right) \left(\eta_i^2 + 2\alpha_{y_i+1} \eta_i\right) \rangle & \text{otherwise} \end{cases}$$

As a result, we replace $Y_i$ with the transformed response

$$\underline{Y}_i \sim \begin{cases} \mathcal{N}\left(-\frac{1}{4(\lambda(\xi_{i2})+\epsilon)} - \alpha_2, \frac{1}{2(\lambda(\xi_{i2})+\epsilon)}\right) & \text{if } Y_i = 1 \\ \mathcal{N}\left(-\frac{1}{4(\lambda(\xi_{iM})+\epsilon)} - \alpha_M, \frac{1}{2(\lambda(\xi_{iM})+\epsilon)}\right) & \text{if } y_i = M \\ \mathcal{N}\left(\frac{\alpha_{y_i}\left(\lambda(\xi_{i,y_i})+\epsilon\right)+\alpha_{y_i+1}\left(\lambda(\xi_{i,y_i+1})+\epsilon\right)}{\left(\lambda(\xi_{i,y_i})+\epsilon\right)+\left(\lambda(\xi_{i,y_i+1})+\epsilon\right)}, \frac{1}{2[\left(\lambda(\xi_{i,y_i})+\epsilon\right)+\left(\lambda(\xi_{i,y_i+1})+\epsilon\right)]}\right) & \text{otherwise} \end{cases}.$$

**Update of $\xi_{im}$:** Following the same argument as in **?**, we have $\xi_{im} = \sqrt{\langle(\eta_i + \alpha_m)^2\rangle}$ to achieve the tightest bound.
**Update of $\alpha_m$:** Given other parameters, we can update the intercepts terms $\infty = \alpha_1 > \alpha_2, \ldots, \alpha_M > \alpha_{M+1} = -\infty$ to maximize the lower bound with some general optimizer (to omit constants). For simplicity, we have let $\lambda(\xi_{im}) \leftarrow \lambda(\xi_{im}) + \epsilon$ in the calculation below:

$$\sum_i \underline{\ell}_i = \sum_{y_i=1}\{-\frac{\alpha_2}{2} - \lambda(\xi_{i,1})(\alpha_2^2 + 2\eta_i\alpha_2)\} + \sum_{y_i=M}\{\frac{\alpha_M}{2} - \lambda(\xi_{iM})(\alpha_M^2 + 2\eta_i\alpha_M)\}$$
$$+\sum_{y_i=2}^{M-1}\left\{\log(1-\exp(-\alpha_{y_i}+\alpha_{y_i+1})) + \frac{\alpha_{y_i}}{2} - \frac{\alpha_{y_i+1}}{2} - \lambda(\xi_{i,y_i})(\alpha_{y_i}^2 + 2\eta_i\alpha_{y_i}) - \lambda(\xi_{i,y_i+1})(\alpha_{y_i+1}^2 + 2\eta_i\alpha_{y_i+1})\right\}$$

Let $\Delta_m = \alpha_{m+1} - \alpha_m > 0$. Then, we want to minimize the neg-loglikelihood:

$$A_1\alpha_2^2 + B_1\alpha_2 + \sum_{m=2}^{M}\{B_k(\alpha_2 + \sum_{l=2}^{m-1}\Delta_l) + A_k(\alpha_1 + \sum_{l}^{m-1}\Delta_l)^2\} + \sum_{m=2}^{M-1} N_m\{\log(1-\exp(-\Delta_m))\}$$

where for $m = 1, \ldots, K$, we define $n_m = |\{i : y_i = m\}|$ and

$$A_m = -\sum_{y_i=m}\lambda(\xi_{im}) - \sum_{y_i=(m-1)}\lambda(\xi_{im}),$$
$$B_m = \frac{n_m - n_{m-1}}{2} - 2\sum_{y_i=m-1}\lambda(\xi_{i,y_i+1})\eta_i - 2\sum_{y_i=k}\lambda(\xi_{i,y_i})\eta_i.$$

We can solve for $\alpha_2$ and $\Delta_2$, …, $\Delta_{M-1}$ under the constraint that $\Delta_m > 0$.

#### C.4. Multi-class logistic regression

A third common response type is the multi-label response. SPEAR models this response type with a multi-class logistic regression problem:

$$\ell_i = \sum_{m=1}^{M}\mathbb{1}(Y_i = m)\log P_m(x_i|\theta,\alpha),$$

where $P_m(x_i|\theta,\alpha) = \frac{\exp(\theta_m^T x_i + \alpha_m)}{\sum_{m'}\exp(\theta_{m'}^T x_i + \alpha_{m'})}$. As a result, when $Y_i = m$, we have

$$\ell_i = \theta_m^T x_i + \alpha_m - \log(\sum_{m'=1}^{M}\exp(\theta_{m'}^T x_i + \alpha_{m'})).$$

As before, to have a closed form integral of the variational distribution, we are going to upper bound the log of the exponential sums with a quadratic function and thus provides a lower bound the log-likelihood. Generalization of **?** to multi-class logistic regression has been studied before in **?**:

$$
\begin{aligned}
\log \sum_{m=1}^{M} \exp(\eta_{im}) &= \zeta_i + \log \sum_{m=1}^{M} \exp(\eta_{im} - \zeta_i) \\
&\le \zeta_i + \sum_{m=1}^{M} \log(1 + \exp(\eta_{im} - \zeta_i)) \\
&\le \zeta_i + \sum_{m=1}^{M} \left\{ \frac{\eta_{im} - \zeta_i - \xi_{im}}{2} + \lambda(\zeta_m)((\eta_{im} - \zeta_i)^2 - \xi_{im}^2) + \epsilon \eta_{im}^2 + \log(1 + \exp(\xi_{im})) \right\}
\end{aligned}
$$

The optimal expressions for $\xi_{im}$ and $\xi$ take the following bound:

$$
\begin{aligned}
\xi_{im}^2 &= \langle (\eta_{im} - \zeta_i)^2 \rangle = \langle \eta_{im}^2 \rangle + \zeta_i^2 - 2\zeta_i \langle \eta_{im} \rangle \\
\zeta_i &= \frac{\frac{1}{2}(\frac{M}{2} - 1) + \sum_{m=1}^{M} (\lambda(\xi_{im}) + \epsilon) \langle \eta_{im} \rangle}{\sum_{m=1}^{M} (\lambda(\xi_{im}) + \epsilon)}
\end{aligned}
$$

As a result, given $\xi_{im}$, $\zeta_i$ and the intercepts $\alpha_m$, we can consider a normal response

$$
\underline{Y}_{im} \sim \mathcal{N}\left(\frac{\mathbb{1}_{Y_{im}=1} - \frac{1}{2} - 2(\lambda(\xi_{im}) + \epsilon)\zeta_i}{2(\lambda(\xi_{im}) + \epsilon)} - \alpha_m, \frac{1}{2(\lambda(\xi_{im}) + \epsilon)}\right),
$$

where the intercepts $\alpha_m$ are estimated as

$$
\alpha_m = \frac{\sum_i \left( (\lambda(\xi_{im}) + \epsilon) \left( \underline{Y}_{im} - \langle \mathbf{u}_i^T \mathbf{B}_j \rangle \right) \right)}{2 \sum_i (\lambda(\xi_{im}) + \epsilon)}.
$$

### D. Additional results on synthetic data

#### D.1. Gaussian Simulation

#### Gaussian Simulation Overview

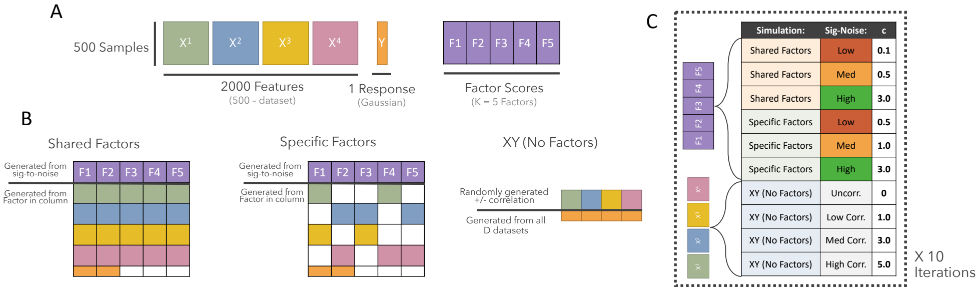

(A) Dimensions of the simulated multi-omic datasets, the Gaussian response of interest and the five simulated factor scores. (B) Representation of the three different types of scenarios: shared factors, specific factors, and no factors. (C) table of each set of parameters tested in the total Gaussian simulation across scenarios, signal-to-noises, and feature correlation.

We simulated synthetic multi-omic data in three distinct scenarios:

1. Shared factors: 5 true factors were generated ($U$), all of which are used to generate $X$ (first 2 factors used to generate $Y$)
2. Specific factors: 5 true factors were generated ($U$), each one randomly influences 2 datasets in $X$ (first 2 factors used to generate $Y$)
3. No factors (XY): $X$ is generated randomly (with varying levels of correlation amongst the features), and $Y$ is randomly generated directly from $X$

In each scenario, $X$ was generated using the `simulate_data(...)` function from the SPEAR package. Parameters were $family = "gaussian"$ (to ensure $Y$ is Gaussian), $N = 500$ (number of training samples), $N.test = 2000$ (number of testing samples), $D = 4$ (number of synthetic multi-omic datasets in $X$ and $X.test$), and $P = 500$ (number of features for each of the $D$ datasets). Each combination of parameters was repeated 10 times across a grid of signal-to-noise ratios (defined below).

The Gaussian scenario when factors were simulated (shared factors and specific factors) was performed in the following manner:

1. Let $U$ represent the simulated factor scores with dimensions $K \times N$ and let $\beta$ represent the simulated loadings matrix with dimensions $P \times K$, where $N$ is the number of simulated samples, $P$ is the total number of simulated analytes, $K$ is the total number of simulated factors, and $c$ is the modulated signal-to-noise coefficient. $c$ takes a grid of values from 0.01 to 5. In particular, we pick c = $(0.5, 1.0, 3.0)$ as low, moderate, and high signal. $\beta$ and $U$ are simulated as follows:

$$\beta_{p,k} = N(0,1) \times c$$

$$U_{k,n} = N(0,1)$$

$$p = 1...P \quad k = 1...K \quad n = 1...N$$

2. Implement sparsity in $\beta$ by nullifying any loadings corresponding to factors that were not designated as influential to $X$. $U_{XY}$ was defined by factors 1 and 2, whereas $U_X$ was defined by all five factors.
3. $X$ was generated in the following manner. Let $E$ represent a matrix with the same dimensions as $X$ ($N \times P$) filled with noise drawn from a normal distribution ($\mu = 0$, $\sigma^2 = 1$).

$$X = \beta * U_{XY} + E \quad E \sim N(0,1)$$

4. $Y$ was generated in a similar fashion. Let $c_2$ represent another sparsity coefficient used to control the spread of $Y$. We kept this parameter consistent at 1 for all our Gaussian simulations. Let 1 represent a vector of all ones utilized to take the row sums, and let $K_Y$ represent the number of factors meant to influence $Y$, which we set to 2.

$$Y = 1^T U_Y \times \sqrt{\frac{c_2}{K_Y}} + E \quad E \sim N(0, 1)$$

When simulating data without underlying factors, $c1$ modifies the correlation amongst the features in $X$ rather than noise in the factors ($U$). In these scenarios, we follow the entire aforementioned procedure with the exception of generating $Y$ as a product of concatenated $X$ and a new vector $B$ of length $P$.

Results for "specific factors" are shown in the main paper. Results for "shared factors" followed the same trends as the "specific factors", only differing in the values of $c1$ for low, moderate, and high signals. Results for "no factors (XY)" are shown in Supplementary Figure 1.

By iterating over a gradient of signal-to-noise ratios, we found a consistent pattern of MOFA performing much worse than SPEAR with high $w$ at a moderate signal-to-noise (1) (Supplemental Fig. 2). This is likely attributed to the lack of supervision in the MOFA model, as further inspection into the SPEAR and MOFA factors (Fig. 2b, Fig. 2c) revealed that MOFA was only able to construct the non-predictive factors in the data.

#### D.2. Ordinal Simulation

To demonstrate SPEAR's ability to model different types of responses, we also carried out additional non-gaussian simulations. Like the Gaussian simulation, five true factors ($U$) were used to construct four multi-omics assays ($X$) and a non-Gaussian response ($Y$) for both training and testing datasets. In the ordinal simulation, the data and response were simulated as follows:

1. Randomly assign 500 training samples and 1000 testing samples a class from $1 - 7$
2. Assign class means ($\mu_1$–$\mu_7$) to be located at the following values: $(-1.5, -1.3, -1, 0, 1, 1.3, 1.5)$. The goal is to generate ordinal class signals that have a nonlinear trend.
3. Generate $U_{\|}$ as the first two factors in $U$: ($U_{\|} = \mu_c + N(500) \times \sqrt{\frac{1}{c}}$). $c$ was modulated to simulate various signal-to-noise ratios. The moderate case shown in the main paper used $c = 11$. The final term is to add noise to the model.
4. Finally, $U_{\perp}$ was generated as the other 3 factors in $U$ randomly ($U_{\perp 1-3} = N(500)$)
5. $X$ was then generated exactly the same way from the Gaussian simulation as described in the manuscript.

We then trained multiple SPEAR models with each treating the response differently (Gaussian, multinomial, and ordinal). In the moderate signal-to-noise, the moderate signal case demonstrated Gaussian SPEAR's difficulty handling the nonlinearity of the class signals in a linear representation (Fig. 3c), further seen in the Gaussian model's predictions (Fig. 3d). SPEAR using multinomial and ordinal responses achieved lower balanced misclassification errors, with the ordinal model outperforming the others. Ordinal SPEAR also extracted predictive signals capable of identifying all seven classes as represented by the correct median probabilities (Fig. 3e).

#### D.3. Multinomial Simulation

The multinomial simulation was carried out similarly to the Gaussian and ordinal simulations, with varying signal levels ($c1$) to simulate low, moderate, and high signal. Results for the moderate signal are shown below, with $c1 = 1$.

Multinomial response $Y$ was simulated in the same manner as the ordinal section above, but with $U_{\|1}$ and $U_{\|2}$ being simulated separately. This is evident in Supplemental Figure 4B below, where certain classes were only distinguishable via either Factor 1 or Factor 2.

### E. Additional results on real data

#### E.1. TCGA-BC dataset background and preprocessing

The TCGA-BC dataset was adapted from Singh et al. (Singh et al., 2019), consisting of 16851 mRNAs, 349 miRNAs, and 9482 CpG methylation sites. Each sample was taken from a primary solid breast cancer tumor that has been classified according to the PAM50 subtype signature, a 50-gene signature. The PAM50 signature is one of the main intrinsic breast cancer signatures used in clinical practice to determine the course of therapy (Wallden et al., 2015; Goldhirsch et al., 2013). Singh et al. showed that even when the 50 genes were removed from the dataset, it was still possible to predict subtype class using the remaining multi-omics features, indicating that there are underlying biological pathways that can distinguish and drive the subtypes.

While most preprocessing was kept as described in Singh et al., including the removal of the PAM50 signature mRNA genes, we additionally removed the least 20% variable features from each assay to reduce the total number of features. We did not exclude the PAM50 genes from the methylation results as the markers are used in gene expression assays. Samples were then split into train ($n_{train} = 379$) and test ($n_{test} = 610$) sets as described in Singh et al. Finally, all three assays were scaled and centered to be normally distributed ($\mu = 0$, $\sigma^2 = 1$).

#### E.2. COVID-19 dataset background and preprocessing

The SARS-CoV-2 (COVID-19) dataset contains multi-omics data from 254 samples from SARS-CoV-2 positive patients and 124 samples from matched healthy subjects from Yapeng et. al. (Su et al., 2020). Samples were taken from one of two timepoints, T1 and T2. At each timepoint, participants were assigned a COVID-19 severity score based on a World Health Organization (WHO) ordinal scale for clinical improvement: uninfected (0), ambulatory without (1) and with activity limitation (2), hospitalized without (3) and with oxygen therapy (4), hospitalized with non-invasive ventilation (5), intubation (6) or ventilation with additional organ support (7) and death (8). These ordinal values were condensed into four classes: healthy (0), mild (1-2), moderate (3-4), and severe (5-7).

MissForest, a non-parametric missing value imputation for mixed-type data was utilized to address missing values in both the proteomics and metabolomics (Stekhoven and Bühlmann, 2012). The metabolite expression values required quantile normalization and log transformation. Both proteomic and metabolomic expression values were then scaled and centered to be normally distributed ($\mu = 0$, $\sigma^2 = 1$).

#### E.3. SPEAR Factor Influence on Prediction

To investigate the predictive influence of each factor generated by SPEAR, we iteratively trained multinomial Lasso classifiers using the *glmnet* R-package with increasing numbers of SPEAR multi-omic factors used as the predictive features. The multi-omic samples were divided into the same training and testing cohorts as described in the SPEAR manuscript. We inspected the balanced misclassification error rate on the training and testing cohorts (Supplementary Fig. 5a, Supplementary Fig. 6a) as well as the coefficient magnitudes for each SPEAR factor for each Lasso model (Supplementary Fig. 5b, Supplementary Fig. 6b).

We found that while Lasso models with fewer SPEAR factors were able to predict well in each dataset, best performance for the prediction of many response classes was found using combinations of many SPEAR factors, suggesting that a singular phenotypes may be driven by multiple complex underlying biological signals.

### F. References

David Wipf,S.N. (2007) A New View of Automatic Relevance Determination. Advances in Neural Information Processing Systems, 20.

Goldhirsch,A. et al. (2013) Personalizing the treatment of women with early breast cancer: highlights of the St Gallen International Expert Consensus on the Primary Therapy of Early Breast Cancer 2013. Ann Oncol, 24, 2206–2223.

Guan,L. (2022) $\ell_1$-norm constrained multi-block sparse canonical correlation analysis via proximal gradient descent. arXiv:2201.05289 [math, stat].

Guan,L. (2021) A smoothed and probabilistic PARAFAC model with covariates. arXiv preprint arXiv:2104.05184.

Guan,L. and Tibshirani,R. (2020) Post model-fitting exploration via a "Next-Door" analysis. Canadian Journal of Statistics, 48, 447–470.

Hastie,T. et al. (2009) The elements of statistical learning: data mining, inference, and prediction Springer.

Singh,A. et al. (2019) DIABLO: an integrative approach for identifying key molecular drivers from multi-omics assays. Bioinformatics, 35, 3055–3062.

Stekhoven,D.J. and Bühlmann,P. (2012) MissForest—non-parametric missing value imputation for mixed-type data. Bioinformatics, 28, 112–118.

Su,Y. et al. (2020) Multi-Omics Resolves a Sharp Disease-State Shift between Mild and Moderate COVID-19. Cell, 183, 1479-1495.e20.

Tian,X. (2020) Prediction error after model search. The Annals of Statistics, 48, 763–784.

Wallden,B. et al. (2015) Development and verification of the PAM50-based Prosigna breast cancer gene signature assay. BMC Medical Genomics, 8, 54.
